# RNA conformational propensities determine cellular activity

**DOI:** 10.1101/2022.12.05.519207

**Authors:** Megan L. Kelly, Rohit Roy, Ainan Geng, Laura R. Ganser, Akanksha Manghrani, Bryan R. Cullen, Ursula Schulze-Gahmen, Daniel Herschlag, Hashim M. Al-Hashimi

## Abstract

Cellular processes are the product of interactions between biomolecules, which associate to form biologically active complexes ^1^. These interactions are mediated by intermolecular contacts, which if disrupted, lead to alterations in cell physiology. Nevertheless, the formation of intermolecular contacts nearly universally requires changes in the conformations of the interacting biomolecules. As a result, binding affinity and cellular activity crucially depend not only on the strength of the contacts, but also on the inherent propensities to form binding-competent conformational states^2,3^. Thus, conformational penalties are ubiquitous in biology and must be known in order to quantitatively model binding energetics for protein and nucleic acid interactions^4,5^. However, conceptual and technological limitations have hindered our ability to dissect and quantitatively measure how conformational propensities impact cellular activity. Here, we systematically altered and determined the propensities for forming the protein-bound conformation of HIV-1 TAR RNA. These propensities quantitatively predicted the binding affinities of TAR to the RNA-binding region of the Tat protein and predicted the extent of HIV-1 Tat-dependent transactivation in cells. Our results establish the role of ensemble-based conformational propensities in cellular activity and reveal an example of a cellular process driven by an exceptionally rare and short-lived RNA conformational state.

---

There is a growing database of nucleic acid and protein structures^6^. However, conformational propensities can only be deduced from conformational ensembles specifying the probability of forming bound states in the absence of binding partners^1^. These bound states can be exceptionally low-populated and short-lived, falling outside detection of standard biophysical methods^7,8^. Moreover, we currently lack approaches for determining conformational ensembles within cells, and conformational propensities may differ within the physiologically relevant cellular environment relative to *in vitro* conditions used to measure them^9,10^. Even under ideal *in vitro* conditions, determining ensembles has been time-consuming^11^, making it difficult to systematically examine how changes in conformational propensities impacts cellular activity. Compounding these limitations are unique challenges in obtaining quantitative and accurate measurements of interactions and their consequences inside cells.

## Thermodynamic model relating conformational propensities to cellular activity

We have taken on this set of challenges, spanning biophysics to cellular function, by quantitatively and systematically examining the role of the transactivation response element (TAR) RNA conformational propensities in Tat-dependent transactivation of the HIV–1 genome (Fig. 1a). TAR is a highly conserved and structured^12^ RNA element located at the 5’ terminal end of the retroviral genome. Transactivation is a multi-step cellular process initiated by binding of the viral protein Tat and the human super elongation complex (SEC) to the active conformation of TAR (Fig. 1a)^13–17^. Productive Tat binding to TAR and cellular transactivation depend on coaxial stacking of the two TAR helices^18^ and the formation of a U23•A27-U38 base triple (Fig. 1b,d), in which bulge residue U23 forms a reverse Hoogsteen base pair with A27 in the upper stem^19–21^. The Tat protein forms several critical contacts with this base-triple^22,23^. Two arginine residues sandwich the base-triple forming base-specific bidentate hydrogen bonds with two guanine residues in the upper stem (Fig. 1c)^20,24^.

**Fig. 1:**
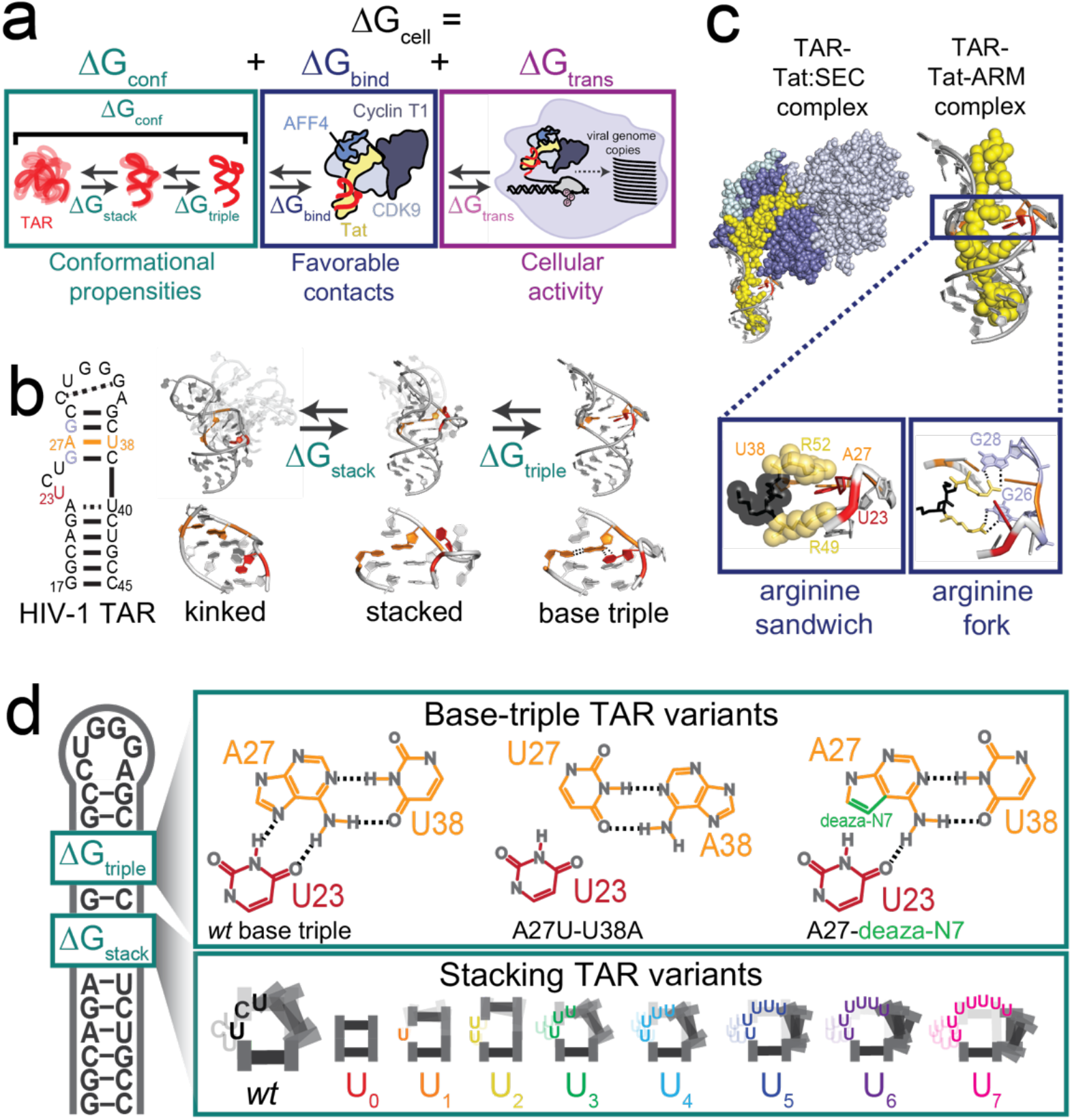
Revealing the role of conformational propensities in HIV-1 Tat-dependent cellular transactivation. **a,** Thermodynamic model of HIV-1 Tat-dependent transactivation. The energetics of cellular transactivation (ΔG_cell_) is decomposed into contributions from conformational penalties to form the base-triple bound TAR conformational state (ΔG_conf_), binding of Tat:SEC to TAR (ΔG_bind_), and the several steps leading to transactivation (ΔG_trans_). **b,** Secondary structure of HIV-1 TAR, FARFAR^27^ models of the bent and stacked ensembles (see Methods), and base triple conformation (PDB entry 6MCE^22^) with close-ups of the base-triple-forming component conformations below. **c,** TAR-Tat:SEC complex (modelled using PDB entries 6CYT and 6MCE)^32^, TAR-Tat-ARM peptide, and critical contacts between TAR and the Tat arginine rich motif (Tat-ARM). Two Tat arginine residues R49 and R52 (in yellow) sandwich U23 in the TAR U23•A27-U38 base triple motif (in red) while also forming hydrogen bonds (dashed lines) via arginine forks with residues G26 and G28 in the TAR upper stem. **d,** Library of TAR variants with two types of mutations which incrementally increase ΔG_stack_ through replacement of the *wt* UCU bulge with increasingly longer uridine bulges (U_0_-U_7_) or increase ΔG_triple_ through replacement of A27-U38 with either U27-A38 or deaza-N7 modified A27. Dotted black lines indicate hydrogen bonds in **c** and **d**.

Prior studies^7^ showed that in the absence of Tat, the free wild-type TAR (*wt*) ensemble has an exceptionally low propensity to form the base-triple bound conformation (Fig. 1b), which was estimated to be energetically disfavored by ΔG > 7 kcal/mol. To quantify how the propensity to form the base-triple impacts Tat-dependent cellular transactivation, we decomposed the energetics of protein-TAR binding (ΔG_prot_) into two independent contributions (Fig. 1a): the conformational propensity to form the base-triple TAR bound state (ΔG_conf_) and favorable binding of the protein to this conformational state (ΔG_bind_) such that

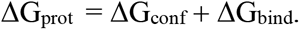

We further decomposed ΔG_conf_ into two contributions (Fig. 1b): stacking of the TAR helices (ΔG_stack_) and formation of the base triple (ΔG_triple_) in the stacked state, again assuming independence,

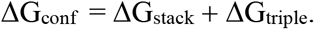

We then examined how mutations that are predicted to alter ΔG_stack_ and/or ΔG_triple_ without affecting contacts between TAR and Tat:SEC, and therefore ΔG_bind_, impact cellular transactivation. This approach directly links RNA conformational preferences to cellular function (Fig 1d). Assuming the TAR variants predominantly bind Tat:SEC in a base-triple conformation, the difference in the measured protein binding energetics, ΔG_prot_, between the reference *wt* TAR and a variant *j* should only depend on the difference in the conformational propensities (ΔΔG_conf_ = ΔΔG_stack_ + ΔΔG_triple_) as follows:

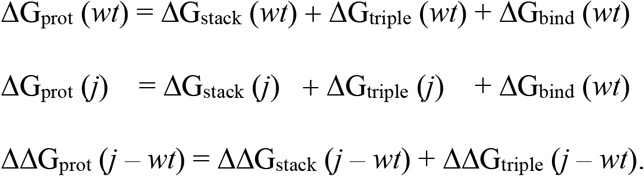

This predictive model allowed us to dissect and learn more about the contribution of conformational propensities to protein binding and cellular function.

The ΔG_stack_ penalty was incrementally increased by increasing the TAR bulge length^18^ from zero to seven nucleotides (U_0_-U_7_) (Fig 1d). Increasing the bulge length leads to increased sampling of kinked conformational states which are non-productive for binding. The ΔG_triple_ penalty was increased for each bulge variant (U0-U7) by destabilizing the U23•A27 base-pair in two ways, by replacing A27-U38 with U27-A38 and by replacing A27 with a deaza-N7-modified adenosine (Fig 1d). Due to topological constraints, the U1 variant is also expected to increase ΔG_triple_ (i.e., render it less favorable) since base triple formation typically requires an additional spacer bulge residue^25,26^.

Assuming the simplest model of ΔG_conf_ (Fig. 1a,b), once a TAR variant becomes stacked, the energy required to form the base triple does not vary with bulge length and is a constant given by ΔG_triple_(*wt*). Our model assumes that all TAR variants are predominantly bound by Tat:SEC in a base-triple conformation (binding to kinked conformations has been detected to form as a minor state with smaller Tat fragments, a point we return to)^7^. Thus, for the base-triple forming variants, the difference between the protein-binding energetics ΔG_prot_ comparing the reference *wt* and a variant *j* is predicted to be equal to the corresponding difference in stacking propensities, DΔG_stack_(*j* – *wt*), which can be measured experimentally^18^,

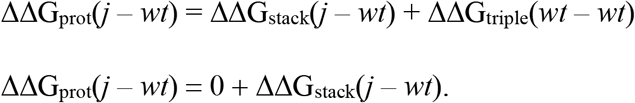

As the model above predicts independent energetic effects from changes in stacking and base triple formation, the base-triple destabilizing variants (denoted *) are predicted to increase ΔG_conf_ relative to their unmodified counterparts by a constant amount (*c*_triple_) for each stacking variant (*j*) where *c*_triple_ is the amount that the base triple is destabilized.

Thus, the following relationships are predicted:

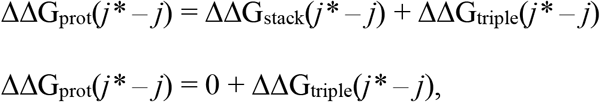

and

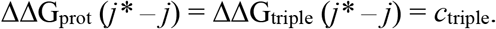

Our model therefore makes strong quantitative predictions: (1) for the bulge variants, ΔΔG_prot_ will vary linearly with ΔΔG_stack_ with a slope of 1 and intercept of 0; (2) for the base-triple destabilized variants (*j*\*), the binding energy of each variant will be weakened relative to its unmodified counterpart (*j*) by a constant amount, *c*_triple_.

## Conformational propensities predict TAR binding to Tat

Before examining how changing ΔG_conf_ impacted Tat-dependent cellular transactivation, we first tested a more direct prediction of the model on *in vitro* binding energetics (ΔG_pep_) of TAR to a 12-amino acid arginine-rich motif (ARM) peptide containing the RNA-binding region of Tat (Tat-ARM peptide) (Fig. 1c). The standard way to measure RNA stacking would be through determination of the full atomic-resolution experimental ensemble. The measured fractional population of the stacked conformation (*p*_stack_) is then:

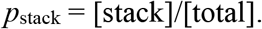

The equilibrium constant defining the stacked state is *K*_stack_:

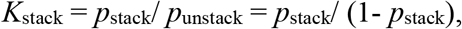

and the free energy of stacking is given by

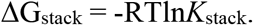

However, determining ensemble typically takes several years per variant^11^. We therefore sought a simpler, but still quantitative approach to determine ΔΔG_stack_ and applied it to our TAR library. We developed such an approach using the NMR chemical shifts of U23-C6 and A22-C8^18^ to measure the fractional population of the stacked conformational state (*p*_stack_) (Fig 2a). This chemical shift perturbation (CSP) approach allowed us to rapidly measure *p*_stack_ and ΔG_stack_ without having to determine complete ensembles.

**Fig. 2:**
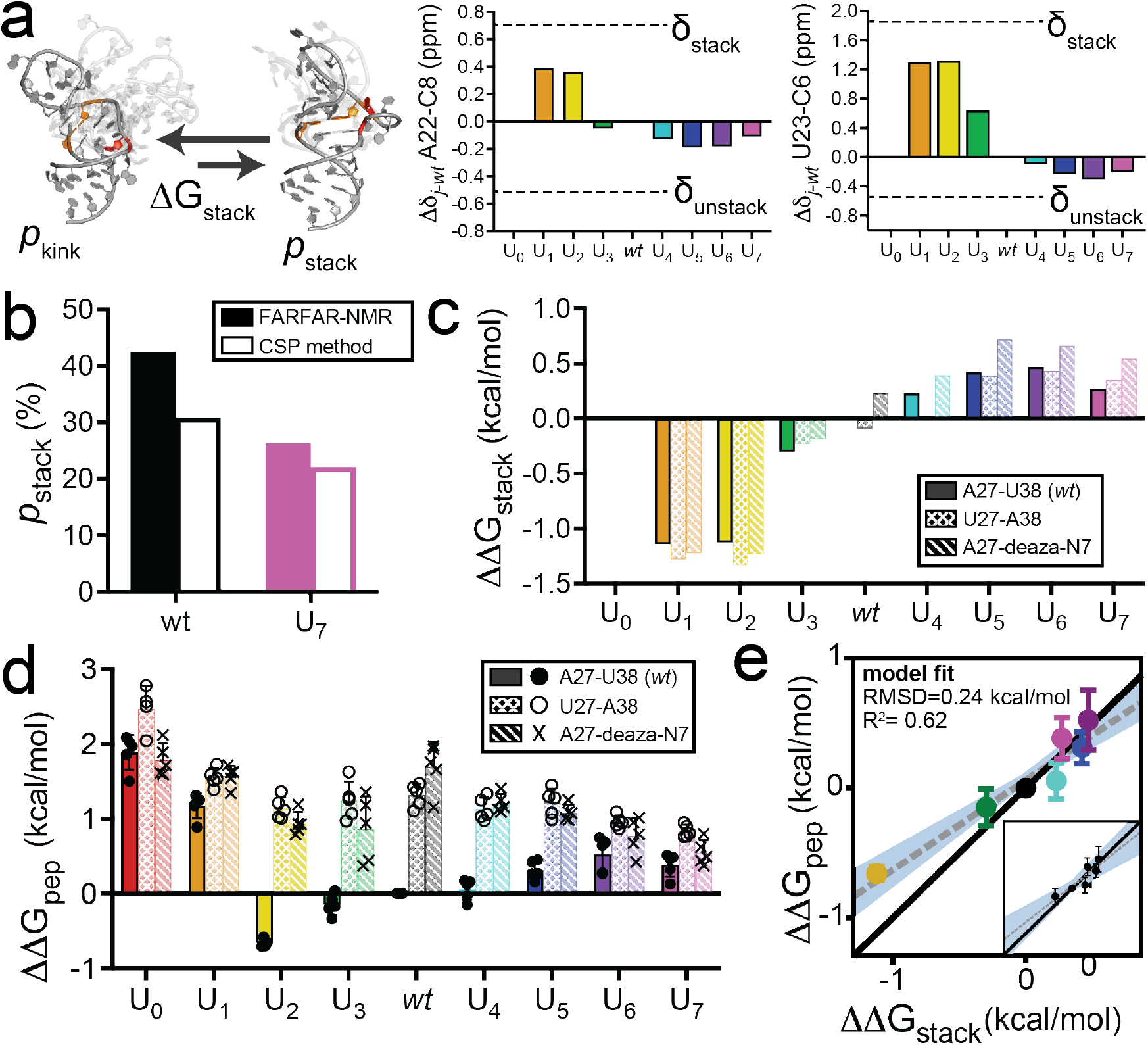
Differences in stacking propensities predict differences in TAR-Tat-ARM binding. **a,** TAR exists in dynamic equilibrium between populations of kinked (*p*_kink_) and stacked (*p*_stack_) inter-helical conformations^18^. Chemical shift perturbations at reporter resonances U23-C6 and A22-C8^18^ are used to measure *p*_stack_. **b,** Comparison of *p*_stack_ deduced using FARFAR-NMR and NMR CSPs (see Methods). **c,** Differences in stacking propensities (ΔΔG_stack_, referenced to *wt*) for the bulge variants with (solid bars) and without the base triple destabilizing A27-deaza-N7 and A27U-U38A mutations (stippled bars) obtained from NMR CSPs. Absolute values of ΔΔG_stack_ are given in Supplementary Table 2. **d,** Differences between the Tat-ARM binding energetics (ΔΔG_pep_, referenced to *wt*) for TAR bulge variants with and without the A27-deaza-N7 and A27U-U38A base-triple destabilizing mutations. Bar height represents the mean, error bars represent standard deviations. **e,** Comparison between ΔΔG_pep_ and ΔΔG_stack_. The black line of slope one indicates predictions from our model. Shown is the fit to this model (RMSD and R^2^), as well as the best fit line (dotted, grey) with the region encompassing the 95% confidence intervals for slope and y-intercept shaded in blue. The inset represents the relationship of the best-fit line (dotted, grey) to the model (black) when U_2_ is excluded from the analysis. The model is now encompassed within the 95% confidence interval values (shaded blue); data points all shown in black for clarity.

We benchmarked the CSP approach by comparing measured ΔΔG_stack_ values with counterparts obtained from full atomic-resolution ensemble models previously developed using NMR-aided Fragment Assembly of RNA with Full-Atom Refinement (FARFAR-NMR)^11,27^. FARFAR-NMR determines RNA conformational ensemble by constraining empirical structure models^27^ with experimental data from NMR spectroscopy^11^. The values of *p*_stack_ were measured for *wt* and U7, the two ensembles with stacked populations in the range of detection for FARFAR-NMR, and were in excellent agreement (Fig 2b, Supplementary Table 1).

Using the CSP approach, we measured *p*_stack_ for all twenty-seven TAR variants used in this study (Extended Data Fig. 1, Supplementary Table 2). The measured *p*_stack_ values across base-triple competent variants varied by 5-fold and corresponds to differences in stacking propensities of ΔΔG_stack_ = 1.6 kcal/mol (Fig. 2c). As expected from entropic considerations and prior results^18^, increasing the bulge length decreased the stacking propensity (Fig. 2c). Unexpectedly, U_7_ did not follow the trend, showing more stacking than U_6_. Additional NMR data provided evidence that U_7_ forms a short uridine-rich helix, which could extrude the bulge and increase the stacking propensity (Extended Data Fig. 2). In agreement with our additivity model, the base-triple destabilizing mutations minimally impacted stacking for all bulge variants across both the A27U-U38A and A27-deaza-N7 variants (r = 0.99 and p < 0.0001 for comparisons of each with the *wt* base triple constructs; Fig. 2c and Extended Data Fig. 1b).

We determined ΔG_pep_ by measuring the binding affinities of the TAR variants to the Tat-ARM peptide using a FRET-based *in vitro* binding assay (Extended Data Fig. 3, Supplementary Table 3)^28^. The range of peptide binding propensities (ΔΔG_pep_) for base-triple competent variants was ~1.2 kcal/mol (Fig. 2d), similar to the range measured for ΔΔG_stack_ of 1.6 kcal/mol.

As predicted by our model, the variation in the binding energetics ΔΔG_pep_ across these variants was in excellent agreement with the differences in stacking propensities, ΔΔG_stack_ (Fig. 2c-d). As expected, the binding affinities decreased with increasing bulge length up to U_6_, and U_7_ showed a higher binding affinity relative to U_6_. A strong linear correlation was observed between ΔΔG_pep_ and ΔΔG_stack_ for base-triple forming mutants (Pearson correlation r = 0.96; p = 0.0006) (Fig. 2e). The predictions from our model, without any adjustable parameters, agree with the experimental data with RMSD = 0.24 kcal/mol and R^2^ = 0.62 (Fig. 2e, Supplementary Table 4). However, the model did not predict all the experimental data within the 95% confidence intervals (CI) for the best-fit slope and y-intercept. A statistical analysis shows that the predominant source of deviation is U2, which did not bind Tat-ARM as strongly as predicted by its stacking propensity (Extended Data Fig. 4a-b). Analysis of the U2 conformational ensemble^11^ reveals that the population of base-tripled state relative to the population of stacked states is lower relative to this ratio for *wt*. This difference is most likely due topological restrictions associated with the shorter bulge^29^ (Supplementary Discussion 1, Supplementary Table 5). Without U2, the model predicted the experimental data within the 95% confidence intervals (CI) for the best-fit slope and y-intercept (Fig. 2e inset). Deviations from our quantitative and predictive model helped identify granular features of the stacked states, information that is needed to determine conformational propensities most accurately and to predict binding and cellular activation.

In contrast to the above results that were consistent with our model predictions, the binding energetics of the base-triple destabilized U_2_-U_7_ variants were not weakened relative to their unmodified counterparts by a constant amount, *c*_triple_, as predicted by the model (Fig. 2d). Rather, they bound Tat-ARM with similar affinities (~100 – 500 nM). In addition, the correlation between ΔΔG_pep_ and ΔΔG_stack_ vanished for these variants (Extended Data Fig. 4c). As binding is nearly uniform across the base-triple variants destabilized by up to ~2.0 kcal/mol relative to *wt*, the results suggest that base triple formation is not required for Tat-ARM binding. Presumably, this different binding mode is favored because the energetic penalty to form the base triple exceeds that needed to bind in the kinked state that lacks the base triple^7^ (Supplementary Discussion 2). Indeed, the distinct fluorescence intensities of the Tat-peptide complex with the base-triple destabilized TAR mutants is consistent with a different Tat-peptide binding mode (Extended Data Fig. 5a-b). Because our model was quantitative and predictive, we could determine when it failed for the base triple mutants and verify that a kinked state can also bind the Tat peptide^7^ (Supplementary Discussion 2).

## Conformational propensities predict Tat-dependent cellular transactivation

Next, we examined whether changes in TAR’s propensity to form the stacked, base-triple conformation (ΔG_conf_), due to our conformational propensity-altering mutations, result in corresponding quantitative changes in Tat-dependent cellular transactivation (ΔG_cell_) as predicted by our simple model (Fig 1a). ΔG_cell_ is defined as the Tat contribution to the overall TAR-Tat:SEC binding energetics inside cells (see Methods, “Calculation of ΔΔG_cell_”). According to our model, the probability of transcriptional activation should be proportional to the probability of forming TAR in its active stacked conformation that contains the *wt* base triple (Fig 1a). Assuming that the multi-step cellular process of Tat-dependent transactivation is under thermodynamic control with respect to Tat:SEC binding to TAR (Fig 3a; Extended Fig. 6) and that the empirically adjusted^30^ cellular concentrations of TAR and Tat:SEC are low enough that most molecules are unbound (“subsaturating”) our model predicts that the observed differences in Tat-dependent cellular transactivation (ΔΔG_cell_) for two TAR variants will be equal to the energetic difference with which they bind the Tat-peptide (see Methods),

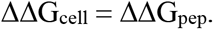

**Fig. 3:**
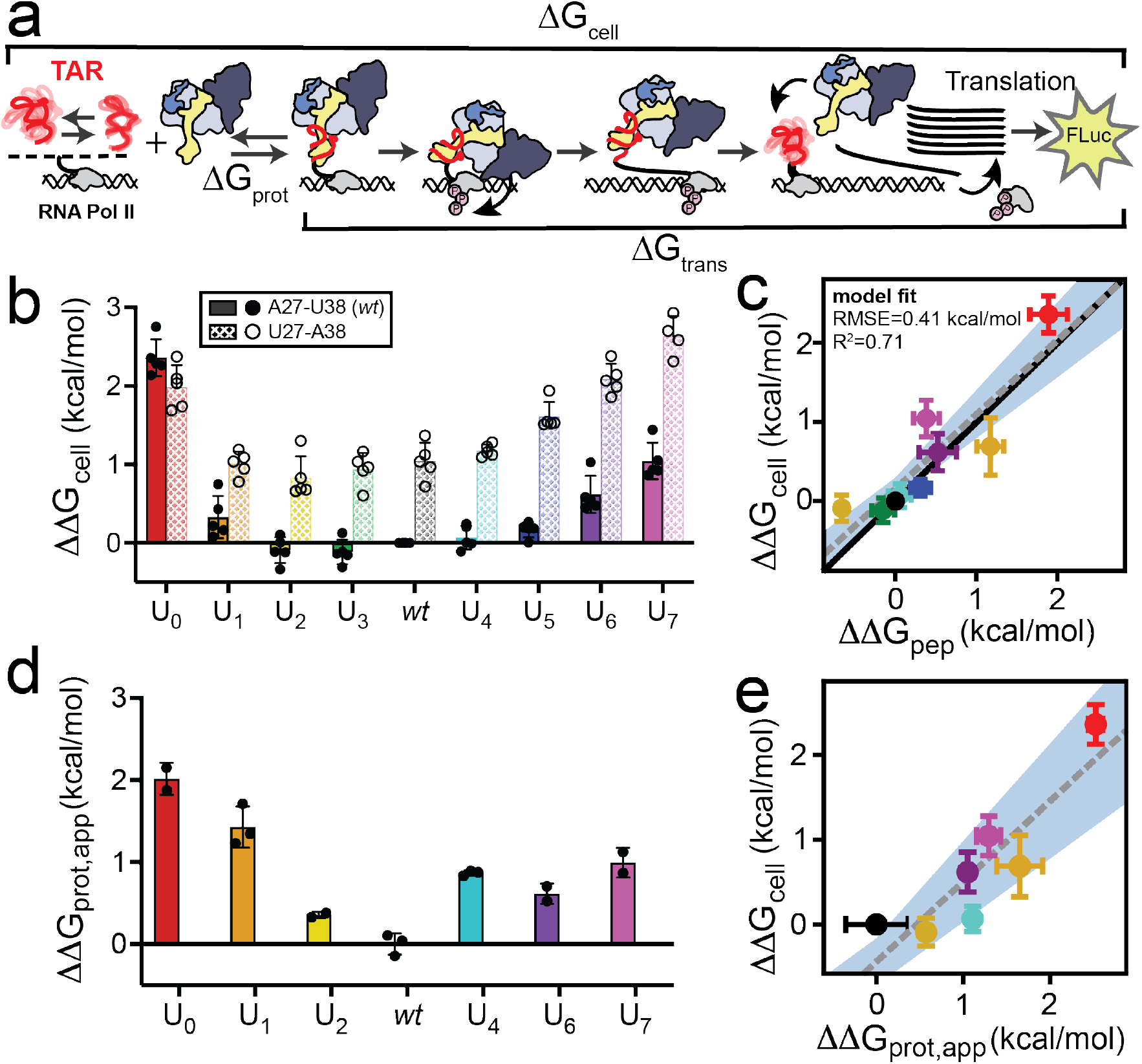
TAR-Tat-ARM binding predicts differences in Tat-dependent cellular transactivation. **a,** Transcriptional activation is a multi-step cellular process which is initiated by binding of the Tat:SEC complex to TAR. The cyclin-dependent kinase 9 (Cdk9) in this complex is then activated, which in turn phosphorylates negative (NELF) and positive (C-terminal domain of RNAP II and Spt5) elongation factors to increase the processivity of RNAPII and activate transcription of the retroviral genome. The energetics of Tat-dependent cellular transactivation (ΔG_cell_) can be decomposed into the conformational penalty of assuming the bound state, mutation sensitive TAR binding to Tat:SEC (ΔG_prot_,app), and contributions from other transactivation steps (ΔG_trans_) assumed to be unaffected by the mutations in our TAR library. b, Differences between cellular transactivation (ΔΔG_cell_, referenced to *wt*) for the bulge variants with (stippled) and without (solid) the base triple destabilizing A27U-U38A mutation. Bar height represents the mean, error bars represent standard deviation. c, Comparison between ΔΔG_cell_ and ΔΔG_pep_ for bulge variants U0-7 *without* the base-triple destabilizing mutation. The black line indicates the prediction from our model. Shown is the fit to this model (RMSD and R^2^), as well as the best-fit line (dotted, grey) with the region encompassing the 95% confidence intervals for slope and y-intercept shaded in blue. d, Differences in the apparent Tat:SEC binding energetics (ΔΔG_prot,app_, referenced to *wt*) for the TAR variants. Bar height represents the mean, error bars represent standard deviation. e, Comparison between ΔΔG_cell_ and ΔΔG_prot,app_ across the TAR variants. The line of best fit is grey and dotted with the region encompassing the 95% confidence intervals for slope and y-intercept shaded in blue.

We used a gene-reporter assay to quantitatively measure ΔG_cell_ for the library of TAR variants^30^, by transiently transfecting (i) the TAR variants driving *Firefly* luciferase (FLuc) expression (ii) Tat under control of a constitutive CMV promoter, and (iii) *Renilla* luciferase (RLuc) also being driven by the CMV promotor to control for transfection efficiency (Fig. 3a). We quantified Tat-dependent cellular transactivation across the different TAR variants from luminescence measurements in the presence and absence of Tat (Extended Data Fig. 6). The cellular concentration of Tat was empirically adjusted by varying the amount of Tat plasmid transfected to avoid saturating TAR (see Methods, Extended Fig. 6). We then converted the measured fold-differences in cellular transactivation into free energy differences (ΔΔG_cell_) (Fig. 3b, Supplementary Table 6).

Remarkably, as predicted by our simple thermodynamic model, we observed quantitative agreement between ΔΔG_pep_ and ΔΔG_cell_ for *wt* and the U_n≥0_ variants (Pearson correlation r = 0.89, p = 0.001) (Fig. 3c). Our experimental data fit the model with an RMSD of 0.41 kcal/mol and an R^2^ value of 0.71 (Fig. 3c, Supplementary Table 4). The model predicts the experimental data to within the 95% CIs of the best-fit slope and y-intercept. These results indicate that the differences in conformational propensities to form the stacked base-triple conformation (ΔΔG_conf_) is similar *in vitro* and in cells and that this propensity determines differences in cellular transactivation (ΔΔG_cell_) (see Fig. 4c).

**Fig. 4:**
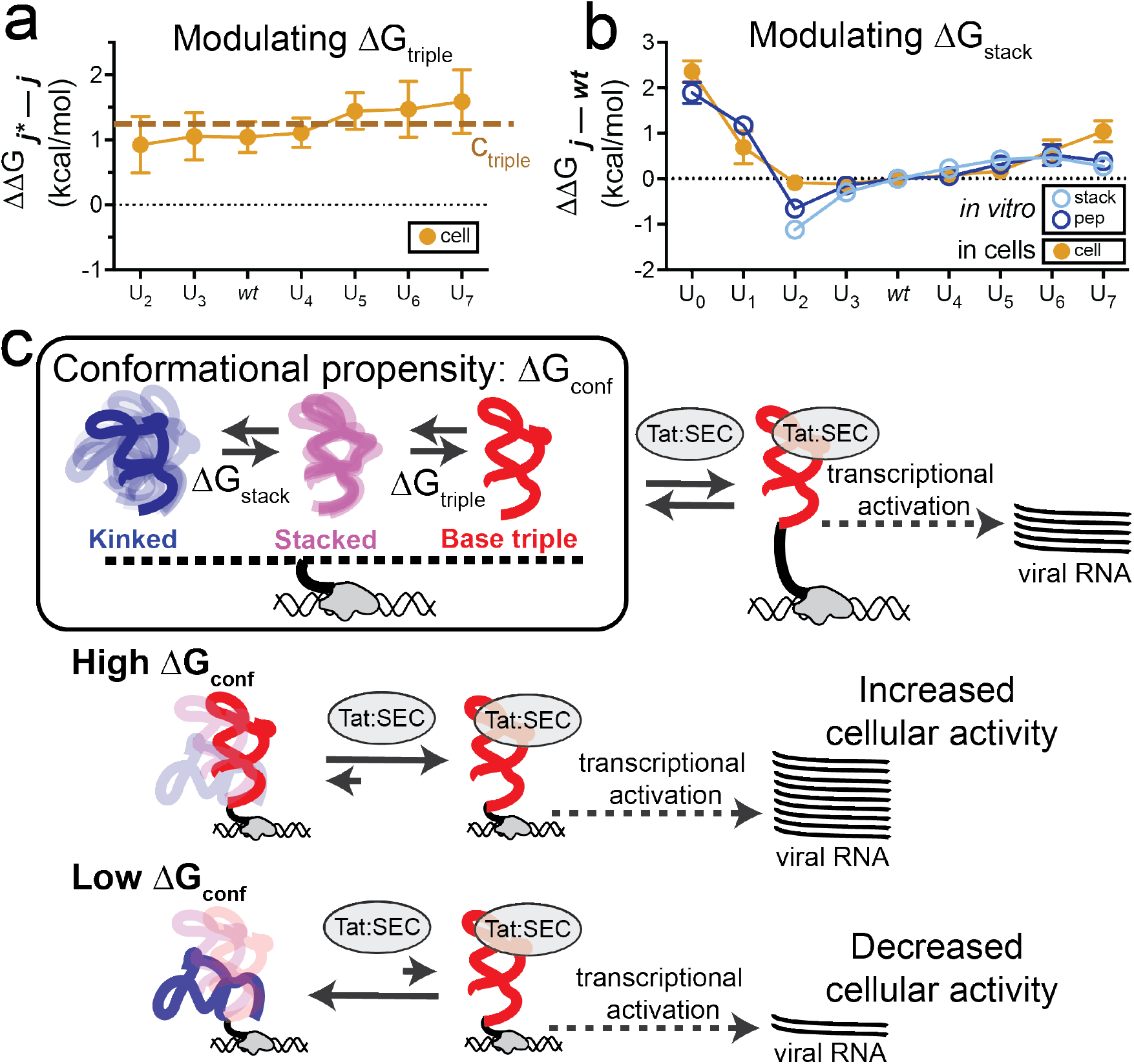
The role of conformational propensities in Tat-dependent cellular transactivation. **a,** Differences in transactivation for *wt* and U_2-7_ variants with the wt-base triple intact (*j*) and with the A27U-U38A base-triple destabilizing mutation (*j*^*^), with dots representing the average value and the errors bars representing the standard deviation. Orange dashed line is the value of ΔΔG_j*-j_ predicted by the model (~1.2 kcal/mol) *), with dots representing the average value and the errors bars representing the standard deviation. **b,** Comparison of ΔΔG_cell_ measured in cells with ΔΔG_pep_, and ΔΔG_stack_ measured *in vitro* for the base-triple forming variants. **c,** Schematic illustrating how conformational propensities shape cellular activity using Tat-dependent transactivation as an example. Increasing or decreasing the conformational propensities to form the RNA conformations bound in the active complex results in corresponding increases or decreases in cellular activity.

To further test that the observed differences in ΔG_cell_ for the TAR variants originate from differences in TAR-Tat:SEC binding as predicted by our model, we semi-quantitatively measured the TAR-Tat:SEC binding *in vitro* for a subset of TAR variants (*wt* and U_1,2,4,6,7_) (Fig. 3d, Extended Data Fig. 7, Supplementary Table 7). We observed excellent agreement (Pearson correlation r = 0.86, p = 0.012) between ΔΔG_prot_,app and ΔΔG_cell_ (Fig. 3e).

Our experiments testing the functional effects of base triple mutations on cellular Tat-dependent transactivation also gave results consistent with our simple model. Specifically, ΔΔG_cell_ for base-triple disrupted mutants was uniformly reduced by a constant (*c*triple) relative to counterparts lacking the base-triple destabilizing mutation (ΔΔG_*j*-j*_ = *c*_triple_ ~ 1.2 kcal/mol; Fig. 4a), in quantitative agreement with predictions from our model (Supplementary Discussion 2). These results suggest that in contrast to Tat-ARM, the Tat:SEC complex preferentially binds these TAR variants in a stacked base-triple like conformation, possibly to ensure contacts also form between the TAR apical loop and the cyclin T1 protein component of Tat:SEC (Supplementary Discussion 2, Extended Data Fig. 5).

Taken together, these results indicate that the difference between the conformational propensities to form the stacked, bound state across the TAR variants are quantitatively the same, within error, *in vitro* and in cells (Fig. 4b). Thus, this fundamental property of the isolated RNA is not altered by differences in *in vitro* versus cellular conditions. We also observed deviations from the model with U2 and U_7_, which indicate directions for further biophysical investigations, and the resulting explanations of these behaviors led to a more complete understanding of this system (Supplementary Discussion 1).

## Conclusions

The impact of mutations on RNA cellular activity is typically interpreted in terms of their effects on intermolecular contacts with proteins or on overall RNA folding^31^. Our study revealed a generally hidden mechanism for tuning RNA cellular activity by altering the propensities to form rare and low-abundance biologically active conformational states (Fig. 4c). Our results demonstrate that conformational propensities depend on both sequence and secondary structure and are needed to quantitatively model RNA cellular activity and understand the evolutionary pressures on RNA sequence and structure. In this regard, it is noteworthy that some of the most common natural variants of HIV-1 TAR, U2 and U3, have the highest stacking propensities and, in some cases, have activities exceeding that of wild-type TAR (Fig. 2c-d, Fig. 3b).

Our findings underscore the importance of going beyond the current paradigm of probing dominant structures *in vitro* and *in vivo* to quantitatively measuring ensembles that describe the propensities to form rare and short-lived biologically active conformational states. By bridging ensembles determined *in vitro* with measurements of cellular activity, the approach established in this study can broadly be applied to quantitatively study the role of conformational propensities in the cellular activities of RNAs and the effect of the cellular environment on these propensities.

## Methods

### Preparation and purification of RNA

Unlabeled wtTAR and TAR mutants (full sequences listed in Supplementary Table 8) for NMR experiments and *in vitro* displacement assays were synthesized with the MerMade 6 DNA/RNA synthesizer (Bioautomation) using standard phosphoramidite chemistry and 2’-hydroxyl deprotection protocols. Samples were purified using 20% (w/v) denaturing polyacrylamide gel electrophoresis (PAGE) with 8M urea and 1X TBE. RNA was excised and then electroeluted (Whatman, GE Healthcare) in 1X TAE buffer. Eluted RNA was concentrated and ethanol precipitated. RNA was then dissolved in water to a concentration of ~50 μM and annealed by heating at 95°C for five minutes and cooling on ice for 1 hour. For NMR experiments, RNA constructs were buffer exchanged using centrifugal concentration (3 kDa molecular weight cutoff, EMD Millipore) into NMR buffer consisting of 15 mM NaH2PO4/Na2HPO4, 25 mM NaCl, 0.1 mM EDTA, 10% (v/v) D_2_O at pH 6.4. For *in vitro* assays, RNA constructs were again buffer exchanged using centrifugal concentration into Tris-HCl assay buffer consisting of 50 mM Tris-HCl, 100 mM NaCl, 0.01% (v/v) Triton X-100 at pH 7.4.

### NMR Experiments

All NMR experiments were performed at 5°C or 25°C, on a Bruker Avance III 700 MHz spectrometer equipped with triple resonance HCN cryogenic probes. Natural abundance RNA samples were exchanged into NMR buffer consisting of 15 mM NaH_2_PO_4_/Na_2_HPO_4_, 25 mM NaCl, 0.1 mM EDTA, 10% (v/v) D_2_O at pH 6.4. Final concentrations ranged from 0.4 to 1.0 mM NMR spectra were processed with NMRPipe^33^ and visualized with SPARKY (version 3.115)^34^.

### Chemical shift perturbation (CSP) measurement of stacking energetics

Stacking populations were calculated from the observed chemical shifts (Δ_obs_) of U23-C6 and A22-C8 for all mutants, measured from 2D HMQC spectra (Extended Data Fig. 2), as described previously^18^. Briefly, Δ_obs_ for a given variant represents a population-weighted average of the fractional populations of the stacked (*p*_stack_) and unstacked (*p*_unstack_) states, such that:

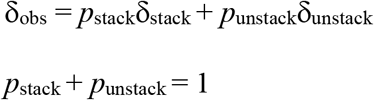

The values of Δ_stack_ (U23-C6) = 143.81 ppm and Δ_stack_ (A22-C8) = 140.19 ppm were obtained from a prior study^18^ based on Mg^2+^ titrations that alter the extent of stacking. The value of dunstack (A22-C8) = 139.00 ppm was obtained from a prior study based on low-salt conditions^35^. The value of dunstack (U23-C6)= 141.50 ppm was obtained based on the most upfield shifted resonance in our set of mutants. The *p*_stack_(A22) and *p*_stack_(U23) values, which agreed within 2-fold for most variants, were averaged to obtain *p*_stack_ for each mutant (Extended Data Fig. 1) and used to determine the ΔG_stack_ value.

### TAR-Tat-ARM peptide binding assay

The fluorescence-based binding assay employed chemically synthesized unlabeled RNA constructs (*wt* and mutants) prepared in-house and a peptide mimic (Genscript) of the Tat RNA binding domain N-AAARKKRRQRRR-C containing the arginine rich motif (ARM), an N-terminal fluorescein label, and a C-terminal Carboxytetramethylrhodamine (TAMRA) label. The peptide is highly flexible when free in solution, allowing the two terminal fluorophores to interact and quench the fluorescent signal^28^. Upon binding to TAR the peptide becomes structured, the two fluorophores are held apart, and fluorescence resonance energy transfers from fluorescein to TAMRA. In this assay, RNA was plated in 1:3 serial dilutions in a 384-well plate and a constant concentration of 20 nM Tat-ARM peptide was used for initial experiments. Both RNA and Tat-ARM peptide were diluted in assay buffer consisting of 50 mM Tris-HCl, 100 mM NaCl, 0.01% (v/v) Triton X-100 at pH 7.4. The plate was left to incubate in the dark for 15 minutes before reading. Fluorescence was measured in triplicate with a CLARIOstar plate reader (BMG Labtech) using a 485 nm excitation wavelength and 590 nm emission wavelength. This assay was repeated five times with distinct samples for each RNA mutant (n = 5).

To ensure the assay accurately reports dissociation constants, the following controls were performed according to Jarmoskaite *et al*, 2020^36^. We first repeated the assay for the tightest binders, *wt* and U_2_, with varying concentrations of the constant component (Tat-ARM peptide) from 2 nM to 100 nM (Extended Data Fig. 3c). Over this 50-fold change in the constant component, we observed at most 2-fold changes in the dissociation constants. These assays were done in triplicate, 1-3 times for each condition. This result provides strong assurance that our assay accurately relates binding constants (is in the so-called “binding regime”^36^). We then tested that the binding reaction reaches equilibrium during the 15-minute incubation. With *wt* TAR, we varied the incubation time from 5 minutes to 2 hours and observed no change in the *K*_d_ (Extended Data Fig. 3d). This was done one time in triplicate, with the same assay plate being read at each timepoint. This constancy provides strong evidence that the reaction has reached equilibrium even at the shortest timepoint for the strongest binder. Baseline fluorescence shifts with increasing incubation times likely result from photobleaching that occurs with each reading of the assay.

The averages and standard deviations of the *K*_d_ values for all variants are reported in Extended Data Fig. 3a and Supplementary Table 3. The. *K*_d_ values were obtained by fitting the binding curves to equation 1 using GraphPad Prism (version 9.3.1),

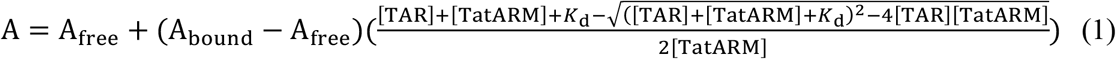

where A is the measured fluorescence; A_free_ is the fluorescence in the absence of TAR-Tat-ARM binding; A_bound_ is the fluorescence at saturated TAR-Tat-ARM binding; *K*_d_ is the measured binding affinity and [TAR] and [TatARM] are the concentrations of TAR and the peptide, respectively.

### Plasmid construction

pcTat^37^, pFLuc-TAR^30^, pcRLuc^30^, and pBC12-CMV^37^ expression plasmids were constructed as described previously. The pFLuc-TAR plasmid was modified to increase expression of FLuc and introduce a BsrgI restriction site for generating TAR mutants. To construct many mutants by inserting pre-annealed oligos with the mutated sequences, we required two restriction sites within 100 bps of each other surrounding the TAR sequence. A HindIII site existed 3’ of TAR, and we then modified the plasmid by making a single base change immediately 5’ of TAR to create a BsrgI site. This was done using two overlapping primers with the desired mutation (Supplementary Table 8). We then eliminated the native BsrgI site in the FLuc gene with a single silent mutation using the same strategy (Supplementary Table 8). This modified plasmid, termed pFLuc-TAR-140, is the wtTAR plasmid in this study. The pFLuc-TAR-140 plasmid was then modified to prepare all the mutants in this study. TAR mutants were introduced at BsrgI and HindIII restriction sites using annealed DNA oligos (Supplementary Table 8) that were designed to have overhangs complimentary to BsrgI and HindIII.

### TAR-Tat dependent trans-activation assay

HeLa cells were maintained in Dulbecco’s modified Eagle medium (DMEM) supplemented with 10% fetal bovine serum and 0.1% gentamicin at 37°C and 5% CO2. Cells were plated to 1.5 x 10^5^ cells per well in 12-well plates 24 hours prior to transfection with polyethylenimine PEI (Polysciences). The primary transfection mixtures contained 125 ng pFLuc-TAR-140 reporter plasmid, 10 ng RLuc control plasmid, +/− 20 ng pcTat expression plasmid, and pBC12-CMV filler DNA plasmid up to a total of 1385 ng total DNA per well. Media was replaced at 24 hours post-transfection. Cells were lysed at 48 hours post-transfection with 250 μL passive lysis buffer (Promega) and incubated 20 minutes at room temperature. FLuc and RLuc activity was measured using a Dual-Luciferase Reporter Assay System (Promega). Assay was repeated in duplicate five times for all mutants on independent days (n = 5).

### TAR-Tat:SEC electrophoretic mobility shift assay

Synthetic TAR RNA was resuspended at 0.1 mg/ml in 20 mM Na HEPES pH 7.3, 100 mM KCl, 3 mM MgCl_2_ and annealed by heating the RNA at 75 °C for 2 min, followed by rapid cooling on ice. Refolded synthetic TAR (nucleotides 17-45) was radioactively labeled with ^32^P-g-ATP using T4-polynucleotide kinase. A 10 μl reaction was prepared with 200 nM TAR, 0.3 mCi ^32^P-g-ATP (7000 Ci/mmol, MP Biomedicals, Sohon, OH), and 10 units of T4-polynucleotide kinase (New England BioLabs, Ipswich, MA) in 70 mM Tris/HCl pH7.6, 10 mM MgCl_2_, 2 mM DTT. After incubating at 37 °C for 1 hour, 25 μl of annealing buffer (20 mM Na HEPES pH 7.3, 100 mM KCl, 3 mM MgCl_2_) were added to the reaction. The mixture was purified twice over Illustra G25 spin columns (GE Healthcare, Piscataway, NJ) to remove free nucleotides. The purified labeled TAR was diluted to 10 nM (3000-5000 cpm/ μl) with annealing buffer for storage and use in EMSAs.

Binding reactions (10 μl) were carried out in 20 mM Na HEPES pH 7.3, 100 mM KCl, 3 mM MgCl_2_, 1 mM DTT, 4% glycerol with 12 units RNasin (Promega, Madison, WI), 10 μg/ml BSA, and 5 μg/ml poly(I:C) (Invitrogen, San Diego, CA). Each reaction contained 50-100 pM labeled TAR RNA. Reactions were incubated at 20 °C for 30 mins and RNA-binding complexes were separated on a pre-run 6% polyacrylamide gel in 0.5x TBE (100 V, 1 hr at 4 °C). Gels were dried, exposed to storage phosphor screens for 12-24 hrs, and imaged on a Typhoon FLA 9000 phosphorimager (GE Healthcare, Piscataway, NJ). The intensity of bands for unshifted and shifted TAR were measured using the program ImageJ (version 1.52)^38^ and the fraction of shifted TAR calculated by dividing the intensity of the shifted TAR by the intensity of the unshifted TAR in the negative control without SEC.

Each EMSA was repeated two to three times on independent days (n = 2-3) and analyzed with GraphPad Prism (version 9.3.1), fitting the EMSA data to equation 2 to calculate apparent *K*_d_ values (Extended Data Fig. 7, Supplementary Table 7):

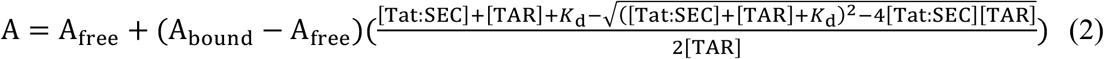

where A is the measured intensity; Afree is the intensity in the absence of Tat:SEC binding; Abound is the fluorescence at saturated TAR-Tat:SEC binding; *K*_d_ is the measured, semi-quantitative binding affinity, and [TAR] and [Tat:SEC] are the concentrations of TAR and the Tat:SEC (Tat:AFF4:P-TEFb) complex, respectively.

The values obtained are referred to as apparent affinities and give apparent free energy values because we were not able to rigorously test whether the tightest binders were subject to titration distortions^36^.

### Calculating *p*_stack_ and interhelical Euler angles for FARFAR-NMR TAR ensembles

The ensemble models for *wt* and variants U_2_ and U_7_, consisting of N = 2000 conformations, used in this study were described previously^11^. In this study, we used these previously derived FARFAR-NMR ensemble models to determine the *p*_stack_ value for *wt* and U_7_. Coaxial stacking of the helices was determined using X3DNA-DSSR (Dissecting the Spatial Structure of RNA) (version 1.9)^39^. DSSR identifies helices as contiguous segments of base-pairs with stacking interactions, regardless of base-pair geometry and backbone connectivity. Conformations in which both upper and lower stems of TAR were identified as part of the same helical stack were assigned to be stacked, all other conformations were assigned to be unstacked (or kinked). The percentage of the conformations in the FARFAR-NMR generated ensembles that were assigned as stacked determined the stacked population (*p*_stack_) value for *wt* and U_7_.

To investigate the deviation of the U2 variant from the model predictions (Supplementary Discussion 1) we analyzed the previously determined *wt* and U2 FARFAR-NMR ensembles^11^ with DSSR (version 1.9)^39^. We used DSSR to assign individual conformations in each ensemble model as stacked or kinked (as above), and as base-triple like or not based on hydrogen bonding between U23 and A27.

### Calculation of ΔΔG_cell_

The gene-reporter assay with a TAR variant (i) quantifies the amount of FLuc mRNA produced during a time interval (t) based on the luminescence signal (*F*(i)) due to the translated FLuc protein:

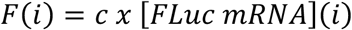

in which *c* is a proportionality constant relating the luminescence signal *F*(i) and the concentration of FLuc mRNA. The FLuc mRNA is produced from Tat-dependent transactivation during which Tat:SEC binds to TAR with association constant *K*_bind,cell_, triggering a cascade of phosphorylation events mediated by the Tat-activated P-TEFb, which modifies positive and negative cellular elongation factors, and results in efficient transcription elongation^17^. This multi-step reaction can be reduced to the following two-step reaction mechanism:

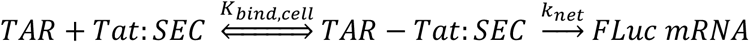

in which *k*_net_ is a kinetic rate constant subsuming all the steps following Tat:SEC binding to TAR leading to FLuc transcription, which is assumed to be equal for all TAR variants. The rate of FLuc mRNA production as a function of time is given by

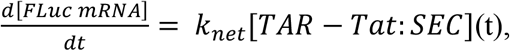

in which [TAR-Tat:SEC](t) is the time-dependent concentration of TAR-Tat:SEC. Assuming that binding is not rate limiting, and *k*net is much slower than the rate of TAR-Tat:SEC dissociation (*k*_net_ ≪ *k*_-1_), we can apply the pre-equilibrium approximation, and obtain an expression for [TAR-Tat:SEC] in terms of *K*_bind,cell_:

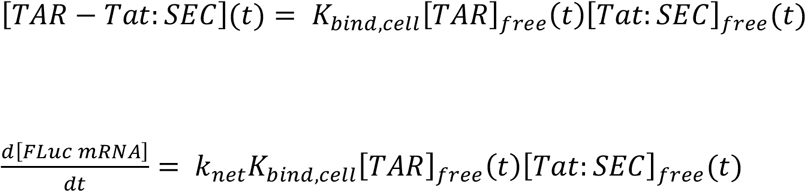

When the total concentrations of [TAR]_total_ and [Tat:SEC]_total_ are much lower than the binding dissociation constant i.e. [*TAR*]_*total*_ and 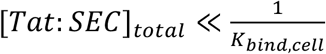, the free concentrations of [TAR]_free_ and [Tat:SEC]_free_ is approximately equal to their total concentrations:

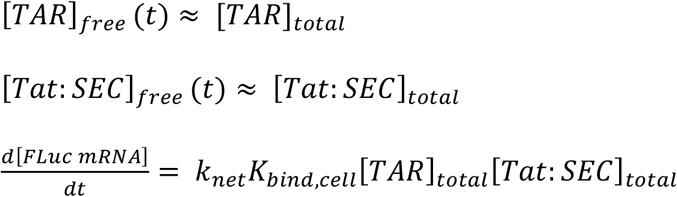

Integrating the above differential equation with respect to time, we obtain the desired expression relating the amount of FLuc mRNA produced over a time interval *τ* and *K*_bind,cell_:

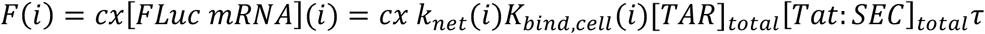

Because only the TAR sequence is altered when comparing results across TAR variants, and since we can control for variations in *k*_net_, [TAR]_total_, and [Tat:SEC]_total_ from one TAR variant to another (see below), the ratio of the FLuc signal (F(i)) measured for a mutant (i) relative to *wt* F(*wt*) will be equal to the fold difference in *K*_bind,cell_:

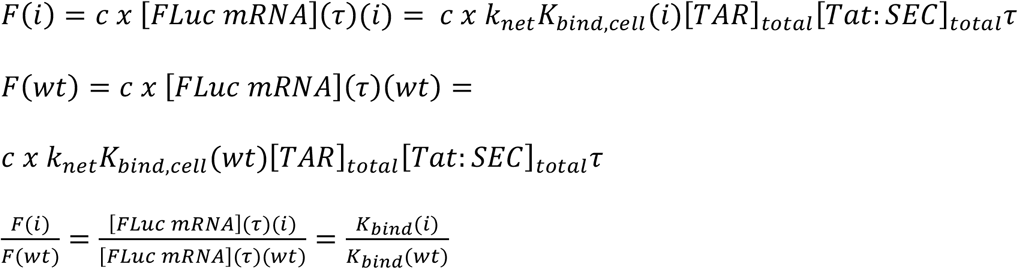

To control for variations in [TAR]_total_ and [Tat:SEC]_total_ due to differences in transfection efficiencies of the DNA plasmids from experiment to experiment, we normalized the FLuc signal by the signal (*R*(i)) obtained for a CMV-driven *Renilla* luciferase (RLuc) reporter plasmid, which is not under the control of the TAR promoter, providing a measure of transfection efficiency and therefore [TAR]_total_ and [Tat:SEC]_total_:

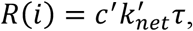

in which c’ is the proportionality constant relating RLuc luminescence to RLuc mRNA, and *k*_net’_ is the net rate of RLuc mRNA production. Provided that the ratio *k*_net_ / *k*_net’_ is constant across TAR variants, the ratio of the RLuc normalized FLuc signal for a mutant (*i*) relative to *wt* will be equal to the fold difference in *K*_bind,cell_:

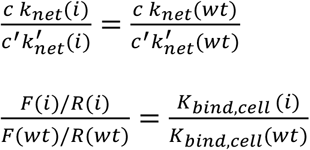

To measure the contribution to *K*_bind,cell_ due to the interaction between TAR and Tat component of the Tat:SEC complex, as well as control for variations in *k*_net_ and *k*_net’_ arising due to differences in the metabolic state of the cells at the time of harvesting, we normalized the FLuc/RLuc signal by the corresponding signal obtained in the absence of Tat:

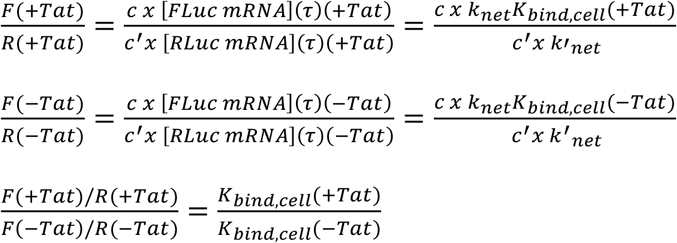

in which *K*_bind,cell_(+Tat) and *K*_bind,cell_(-Tat) are the equilibrium association constants for binding of Tat:SEC and SEC only to TAR in cells, respectively:

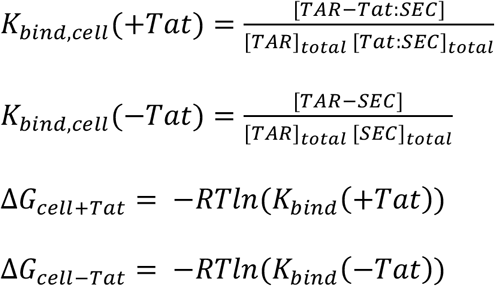

Based on our additivity model, we can decompose the energetics of Tat:SEC binding to TAR into energetic contributions arising from interactions between TAR and Tat (ΔG_Tat_) and between TAR other components of SEC (ΔG_SEC_). The binding energetics with and without Tat are given by:

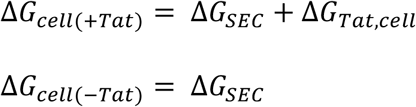

The Tat contribution to the overall TAR-Tat:SEC binding energetics, which we define as the Tat-dependent cellular transactivation (ΔG_cell_) is then given by:

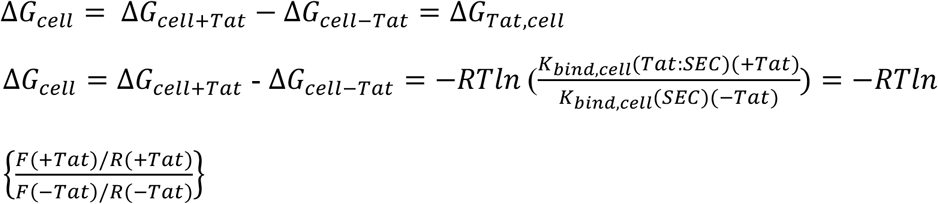

To ensure [*TAR*]_*total*_ and 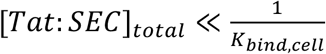, the concentration of the Tat plasmid was empirically adjusted to ensure that we (i) observe differences between *wt* and the mutants^30^ and (ii) fall in the linear proportionality regime in which increasing the Tat plasmid over a 10,000-fold range leads to a corresponding linear increase in the FLuc signal (Extended Fig. 6c-d).

### Statistical comparisons of observed *vs.* predicted ΔΔG values

Statistical analysis was done using GraphPad Prism (version 9.3.1). For all ΔΔG dataset comparisons, three statistical analyses were performed: (1) a Pearson correlation (r) and associated *p* value was computed using a two-tailed t-test; (2) a linear regression to the line of best fit, accounting for the number of replicates and standard deviations, was performed, and the 95% confidence intervals of the best fit slope and y-intercept were calculated; and (3) a least squares regression was used to fit the comparisons to our model with a fixed slope of 1 and fixed y-intercept of 0, which output R^2^ and RMSD values.

### Z-score analysis for ΔΔG_stack_ vs. ΔΔG_pep_

To test the null hypothesis of our model, that the ΔΔG_stack_ values are found in the same distribution as the ΔΔG_pep_ values for each individual mutant, we determined the Z-scores for each variant for the correlation of stacking and peptide binding using,

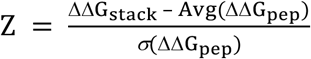

This analysis resulted in the Z-scores reported in Extended Data Fig. 4a. Assuming a normal Gaussian distribution, we used a *p* value cutoff of 0.05 (Corrected for multiple comparisons using the Bonferroni method to 0.0083) to determine our Z-score cutoff for determining outliers, which was Z = 2.4. The distribution of differences between each ΔΔG_pep_ replicate value and its corresponding ΔΔG_stack_ value bulge variants U_2-7_ is shown in Extended Data Fig. 4b.

### Figure design

All figures were designed in Adobe Illustrator 2020 (version 24.1). Images originally created from other sources were exported to Adobe Illustrator to be incorporated into final figures. Bar graphs were plotted in GraphPad Prism (version 9.3.1). Correlation plots were plotted using Python3. Chemical structures were created using MarvinSuite version 19.21.0 (2019) from ChemAxon (https://www.chemaxon.com). Structural models of RNA and proteins were formatted in MacPyMOL (version 1.5.0.4). NMR spectra were processed using NMRPipe^33^, visualized using Sparky (version 3.115) ^34^, and then exported to Adobe Illustrator.

## Data and materials availability

Code written to analyze FARFAR-NMR ensembles can be found on GitHub: https://github.com/alhashimilab/TAR_Thermodynamic_Model/. All other data are available in the main text or supplementary materials. Further information is available from the corresponding authors upon request.

## Acknowledgements

We would like to thank the Duke Magnetic Resonance Spectroscopy Center for nuclear magnetic resonance resources. We would also like to thank John Shin for assistance with statistical analysis. This work was supported by the National Institutes of Health (NIH)/ National Institute of General Medical Sciences (NIGMS) grant R01GM132899 to H.M.A.-H. and D.H., as well as the NIH National Institutes of Allergy and Infectious Disease (NIAID) grants U54 AI150470 to H.M.A.-H., F30 AI143282-01A1 to M.L.K., and 5R21AI156915 to U.S.-G.

## Author contributions

M.L.K, H.M.A., and D.H. conceptualized the study. M.L.K., R.R., A.G., L.G., U.S.G designed the experiments and collected the data. M.L.K performed the NMR experiments and analyzed the data. R.R. performed FARFAR-NMR analysis. M.L.K., A.G., and R.R. performed *in vitro* Tat-ARM peptide binding experiments, and M.L.K. analyzed the data. M.L.K., A.G., and L.G. performed the cellular transactivation experiments, and M.L.K. analyzed the data. M.L.K., R.R., and A.M. conceptualized the methodology for calculating free energy of cellular transactivation. U.S.G. performed the *in vitro* TAR-Tat:SEC binding assays and analyzed the data. M.L.K. created the figures. H.M.A. and D.H. acquired funding. H.M.A, D.H., and B.R.C. supervised the study. M.L.K., H.M.A., and D.H. wrote the manuscript with input from the remaining authors.

## Additional Information

Supplementary information is available for this paper. Correspondence and requests for material should be addressed to: Hashim M. Al-Hashimi; Daniel Herschlag; and Ursula Schulze-Gahmen.

---

**Extended Data Figure 1.**
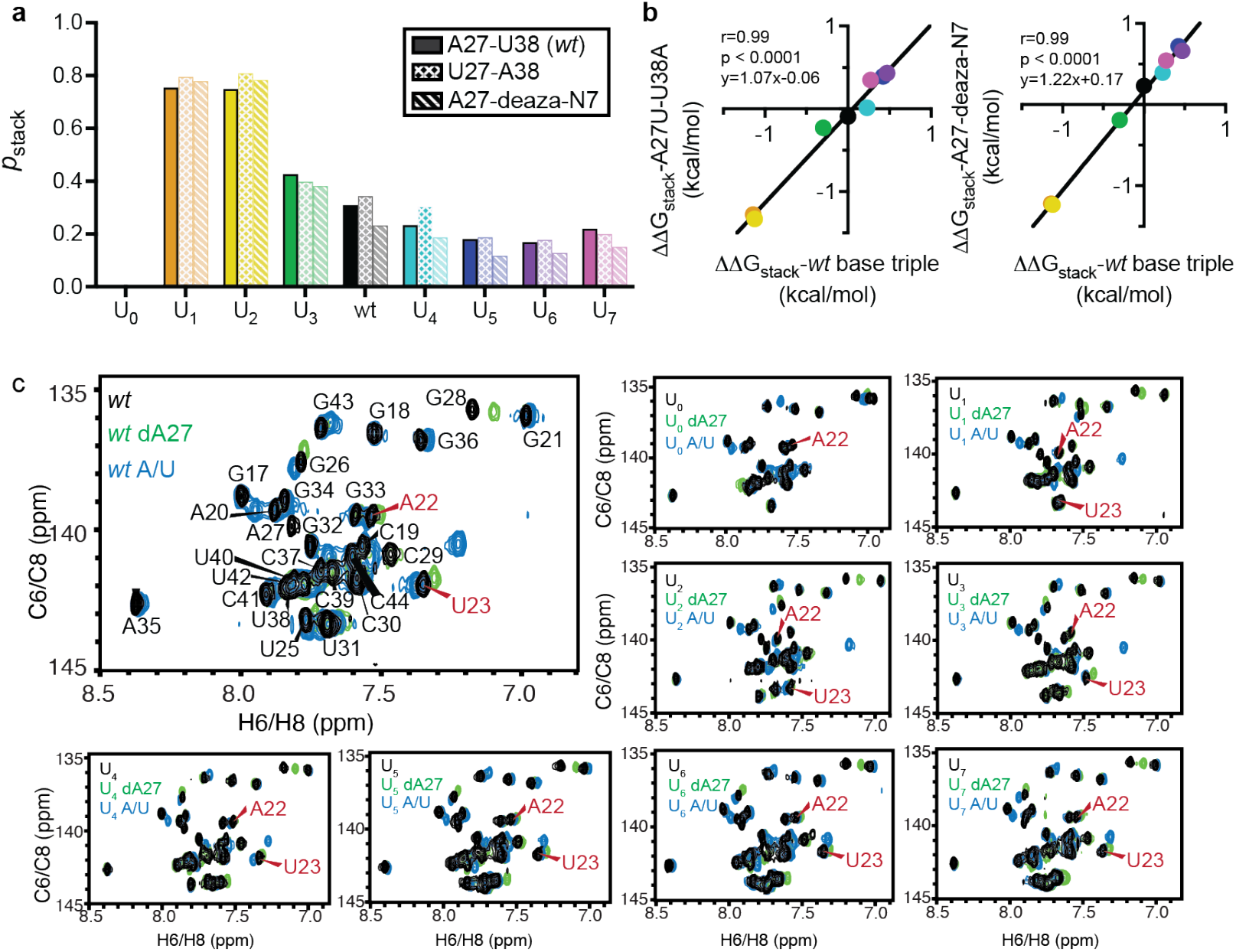
Measurement of *p*_stack_ by 2D aromatic [^13^C, ^1^H] SOFAST-HMQC^40^. **a**, *p*_stack_ (see Methods) for all TAR mutants U0-7 and *wt* with and without the A27U-U38A and A27-deaza-N7 base-triple disrupting mutations. **b,** Correlations between stacked populations for the wt base triple mutants versus A27U-U38A (left) and A27-deaza-N7 (right) mutants. Pearson correlation (r) and line of best fit is shown. **c,** Sets of overlayed spectra for *wt* and all bulge mutants U0-7. For each, the *wt* base triple construct is black and the base-triple disrupting mutants are overlayed, A27U-U38A in blue and A27-deaza-N7 in green. The *wt* spectrum is fully assigned, for the bulge constructs the stacking reporter residues A22 and U23 are indicated.

**Extended Data Figure 2.**
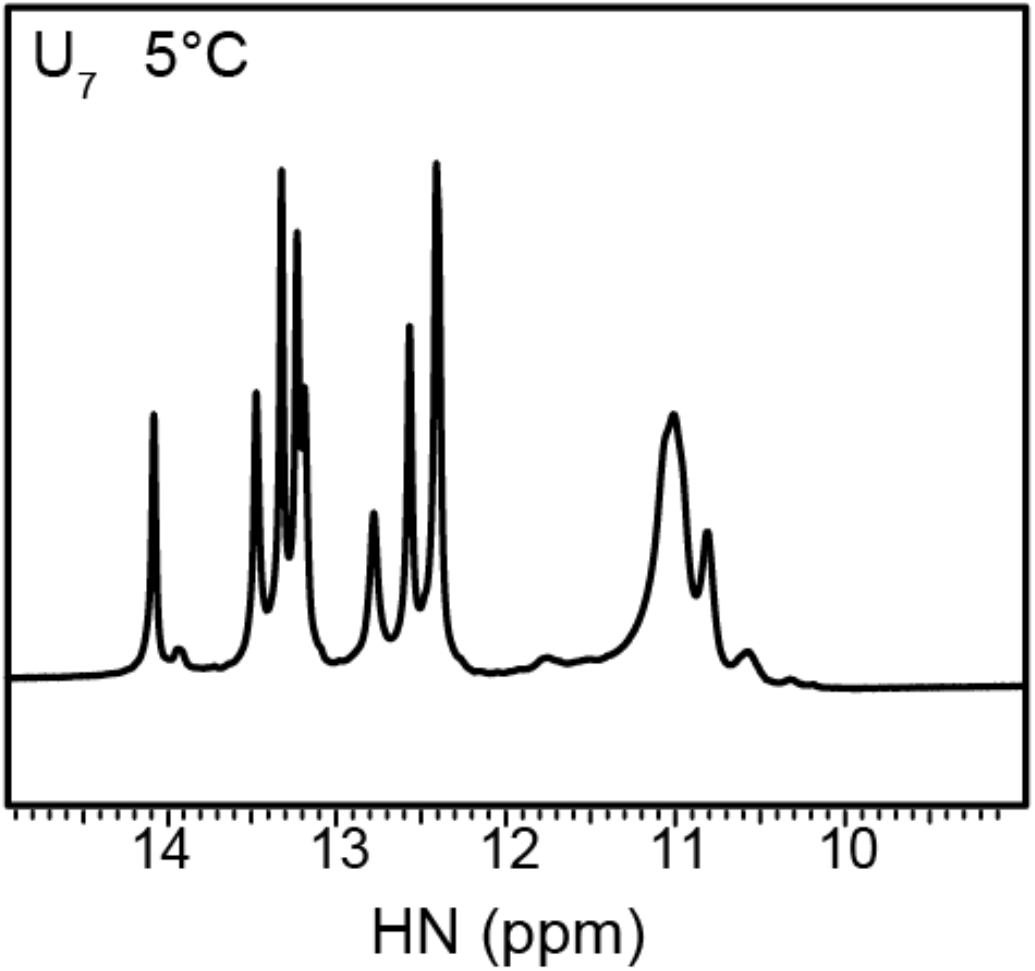
NMR evidence for U-U wobbles in the U_7_ TAR variant. The ^1^H 1D imino NMR spectrum of the U_7_ variant shows resonances in the 10-12 ppm region, suggesting the U-rich bulge might transiently form a short helix comprised of U-U wobble mismatches which could in turn promote stacking of the TAR helices.

**Extended Data Figure 3.**
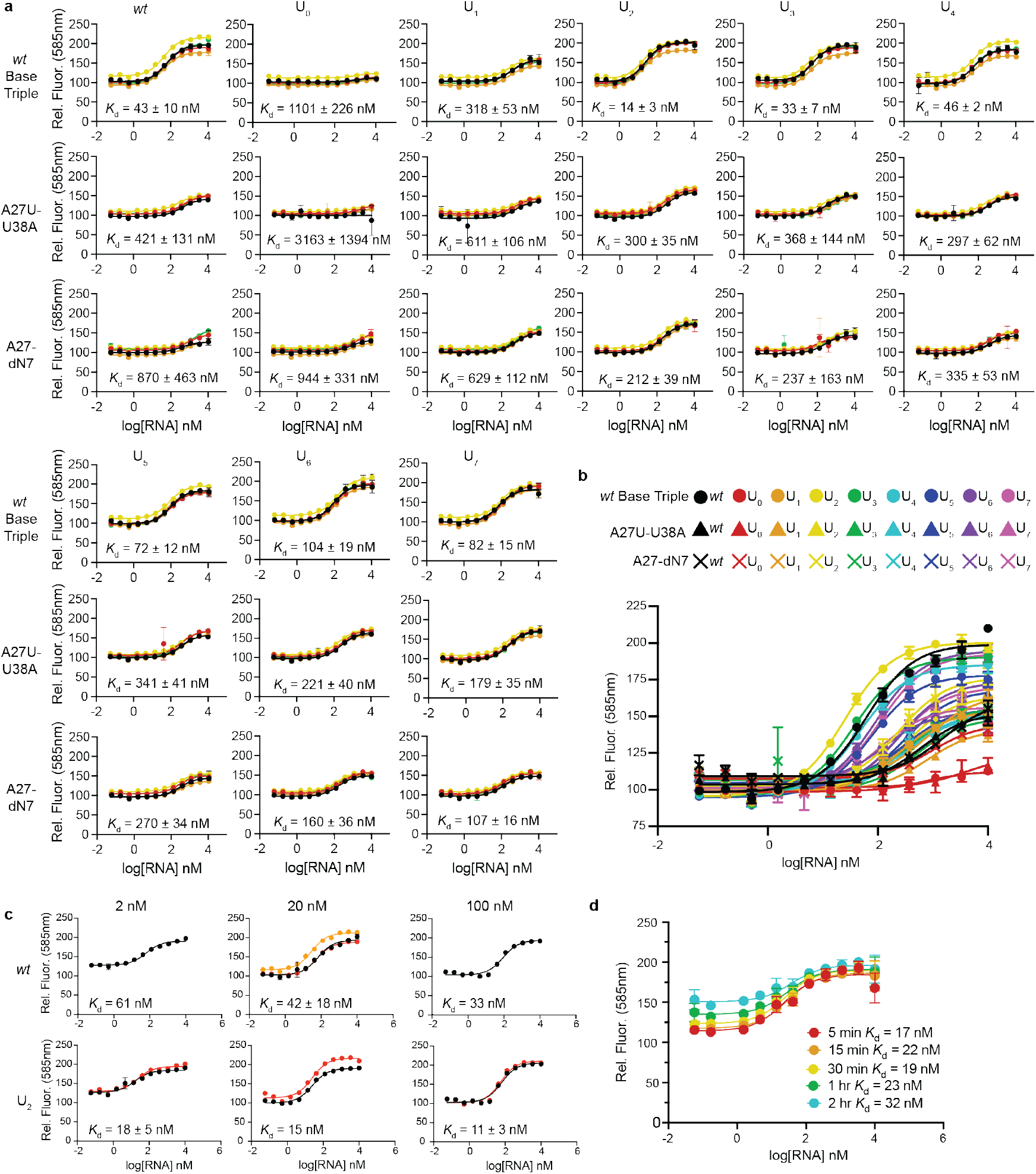
TAR-Tat-ARM peptide binding assay. **a,** Binding curves for individual TAR variants, with all five replicates overlayed (black: replicate 1, red: replicate 2, orange: replicate 3, yellow: replicate 4, green: replicate 5). Data were fit to equation 1, and average *K*_d_ values over the five replicates are displayed for each mutant.**b,** One replicate (replicate 5) of representative fluorescence binding curves for all TAR mutants overlayed. **c,** Observed dissociation constants do not change as the concentration of the constant component (Tat-ARM peptide) is varied, as expected for accurate *K*_d_ measurements^36^. Dissociation constants were measured for *wt* and U2 at multiple concentrations of Tat-ARM peptide, varying 50-fold. The dissociation constants for *wt* and U2 remain vary < 2-fold over this range. **d,** Observed dissociation constants do not change as the equilibration time is varied, as expected for accurate *K*_d_ measurements^36^. Shown are *K*_d_ measurements for *wt* at varying timepoints to demonstrate the reaction has reached equilibrium. The *K*_d_ value does not decrease with increasing incubation times, indicating the reaction has reached equilibrium at the lowest timepoint. The same assay plate was read at each time point, creating a photobleaching effect at each subsequent timepoint, which is evident in the increasing baseline values.

**Extended Data Figure 4.**
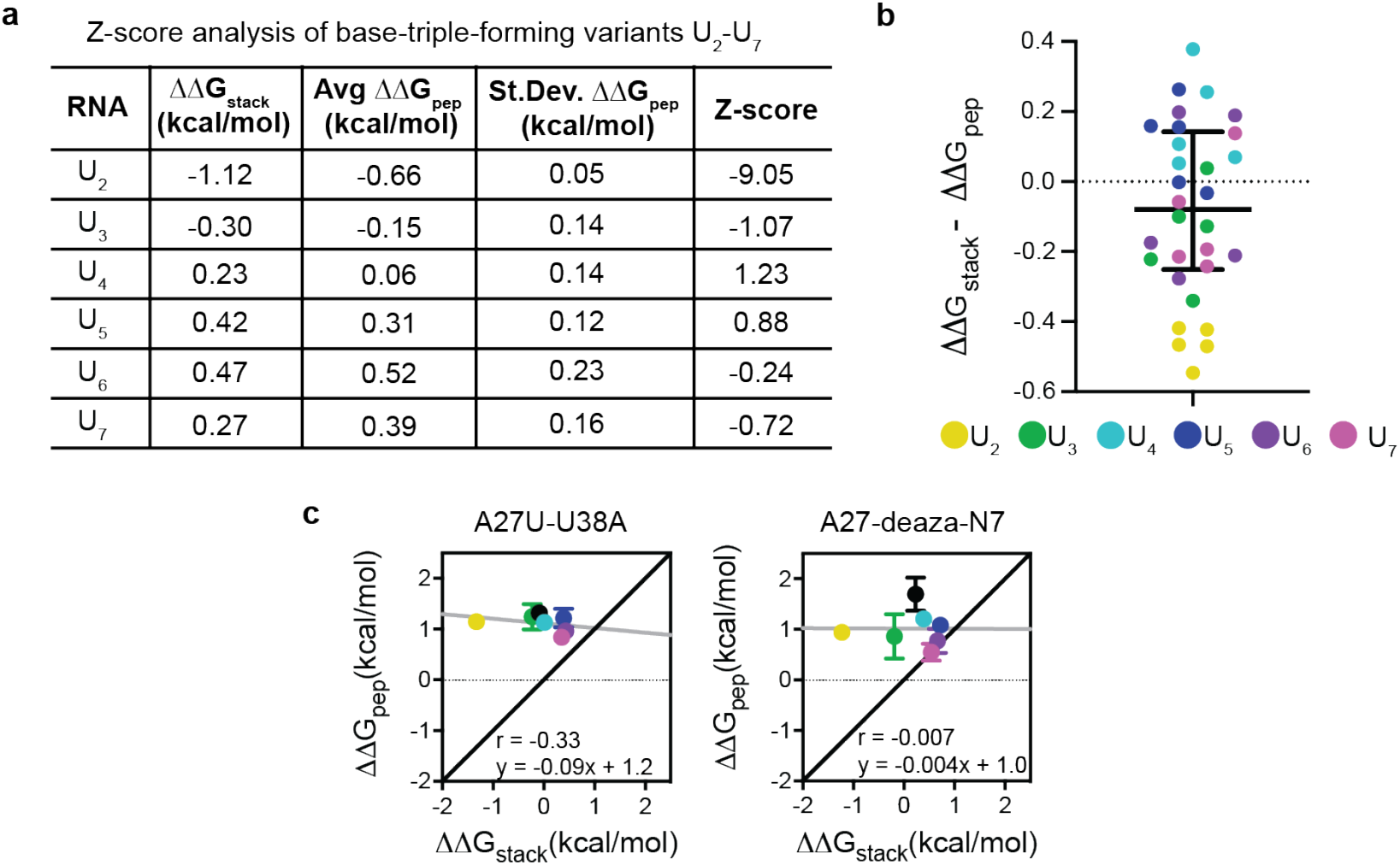
Stacking and peptide binding energetics for *wt* and U_2-7_. **a,** Z-score analysis for base-triple forming variants (see Methods) with the null hypothesis being that the values for ΔΔG_pep_ and ΔΔG_stack_ are from the same distribution for each variant. A Z-score limit of 2.4 rejects the null hypothesis with Bonferroni corrected *p* value of 0.083. U_2_ is the only variant outside this limit. **b,** Distribution of ΔΔG_pep_ deviations from ΔΔG_stack_ per variant (color legend in panel), showing deviations for U_2_. The median and interquartile range shown in black lines. **c,** ΔΔG_pep_ *versus* ΔΔG_stack_ for base-triple destabilized mutants, A27U-U38A (left) and A27-deaza-N7 (right), correlates poorly (Pearson correlation shown). Grey lines indicate the best fit (equation shown), and black lines indicate slope of 1, which is the prediction of the model in the absence of the base triple disrupting mutations.

**Extended Data Figure 5.**
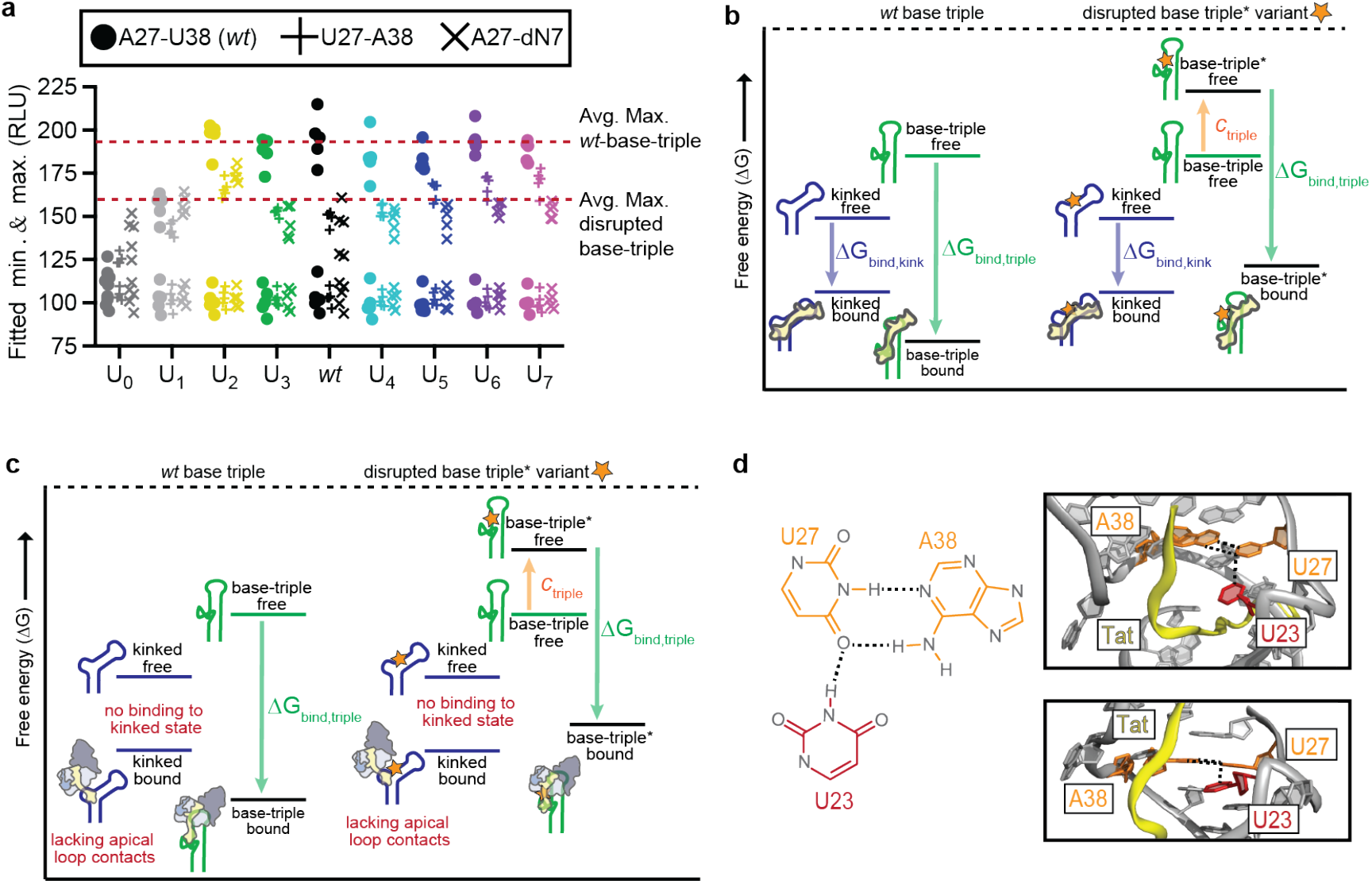
Energetics of base-triple disruption in Tat-ARM binding and cellular transactivation. **a,** Changes in fluorescence upon peptide binding is greater for base-triple competent variants than for base-triple disrupted variants. Shown are the fitted minimum and maximum fluorescence values (from equation 1, see Methods) from the TAR-Tat-ARM peptide binding assay. Red dotted lines indicate average maximum values for the base-triple competent variants (190) and base-triple disrupted variants (155). U_0-1_ are shown in grey as they are unable to form the base-triple. **b,** Energy diagram of Tat-ARM peptide binding to base triple competent and base-triple disrupted variants. The peptide can bind a bulge-independent kinked TAR conformation lower in energy than the base-triple disrupted stacked conformation. **c**, Energy diagram of Tat:SEC binding to TAR in the cellular context. The favorable interactions between Cyclin T1 and the TAR apical loop are unable to form in the kinked state of TAR, and so each base-triple disrupted variant is destabilized by the same amount (*c*_triple_) and binds its non-base triple stacked state (demarcated with an asterisk*). **d,** Proposed model for an alternative sheared base-triple conformation in the A27U-U38A base-triple disrupting mutants with hydrogen bonds shown as black dashed line (left). Two views of the 3D structural model for the alternative sheared base-triple conformation obtained by replacing A27 with U and U38 with A in the PDBID:6MCE^22^ U_2_ TAR structure (right).

**Extended Data Figure 6.**
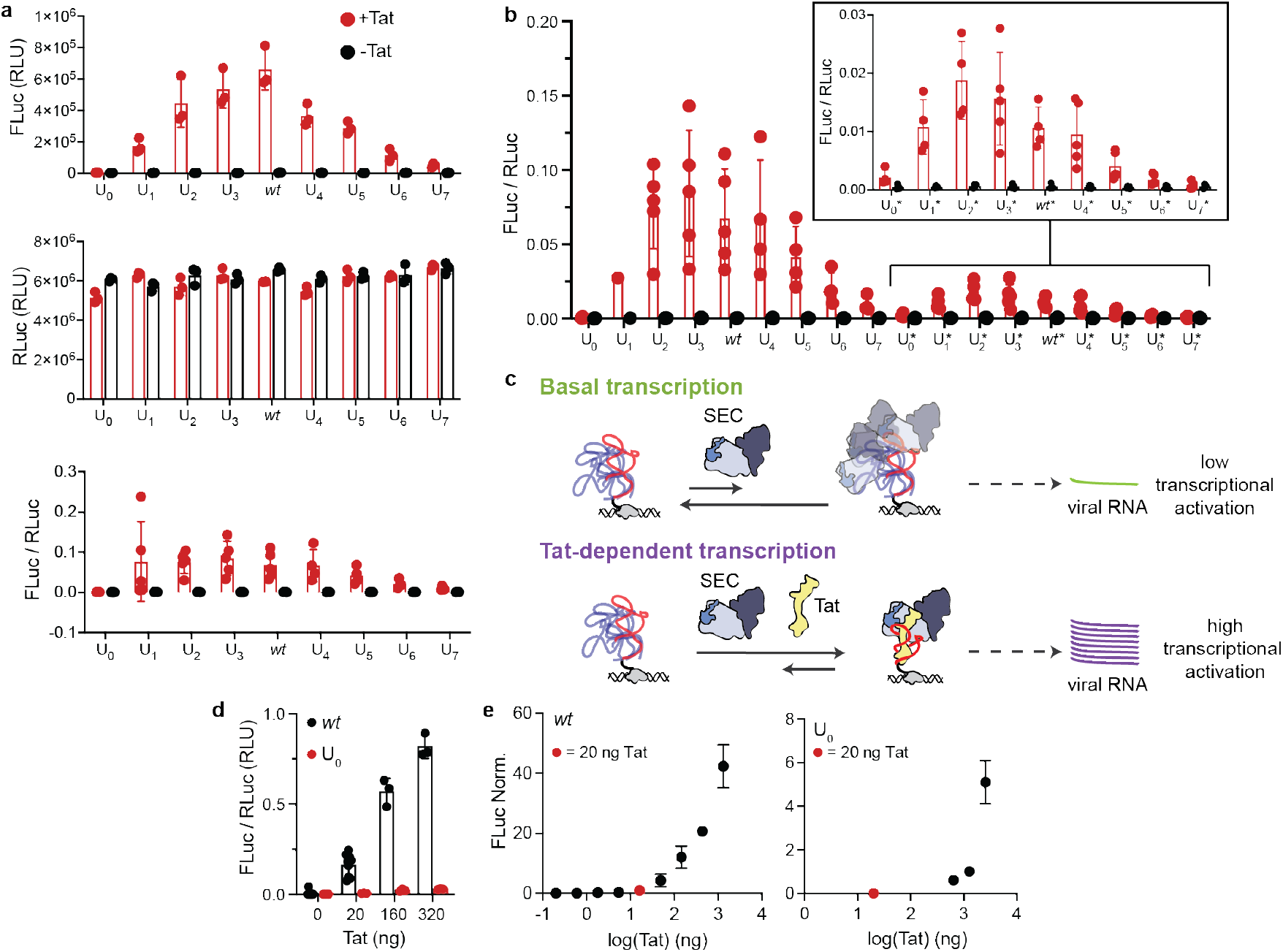
Cellular transactivation assay. **a,** Representative example of luminescence data for one biological replicate of U_0-7_ and *wt* (5 technical replicates). Shown are luminescence values for Firefly luciferase, reporting on transactivation (top), luminescence values for *Renilla* luciferase under control of a CMV promoter, used as a control for transfection (middle), and the ratio FLuc/RLuc to normalize for differences arising from transient transfection (bottom). **b,** Representative example of FLuc/RLuc data for all TAR mutants including base-triple disrupting mutants. Mutants labelled with (*) indicate the A27U/U38A base-triple disrupting mutation. In all graphs, red data are values when Tat is co-transfected and black data are values without Tat, representing Tat-independent baseline activity. **c,** Model of Tat-dependent versus Tat-independent transactivation energetics in cells. (Top) The observed level of basal transcription is likely due to many nonspecific binding interactions of the preformed SEC complex to TAR, which does not alter the conformational propensities of the TAR ensemble and has a low probability of achieving an active bound conformation leading to transactivation and transcription. (Bottom) In Tat-dependent transactivation, the presence of Tat increases the binding affinity to form the active bound state, leading to higher levels of transactivation and transcription. **d,** Tat plasmid titration. In this experiment, the concentration of Tat was varied for *wt* (black), one of the most transactivating constructs, and U_0_ (red), one of the least transactivating constructs. We see that for both *wt* and U_0_, the level of transactivation (FLuc/RLuc) increases with an increase in Tat, indicating that the reaction is not saturated at the level of Tat we are using (20 ng). **e,** Larger scale Tat plasmid titrations for *wt* and U0 covering four orders of magnitude, with the y-axis being FLuc signal normalized to the average FLuc value measured for *wt* at 20 ng Tat. Again, for both mutants, the level of transactivation continually increases with an increase of Tat plasmid; the value we use in our assays (20 ng, red dot) is at the low end of this spectrum.

**Extended Data Figure 7.**
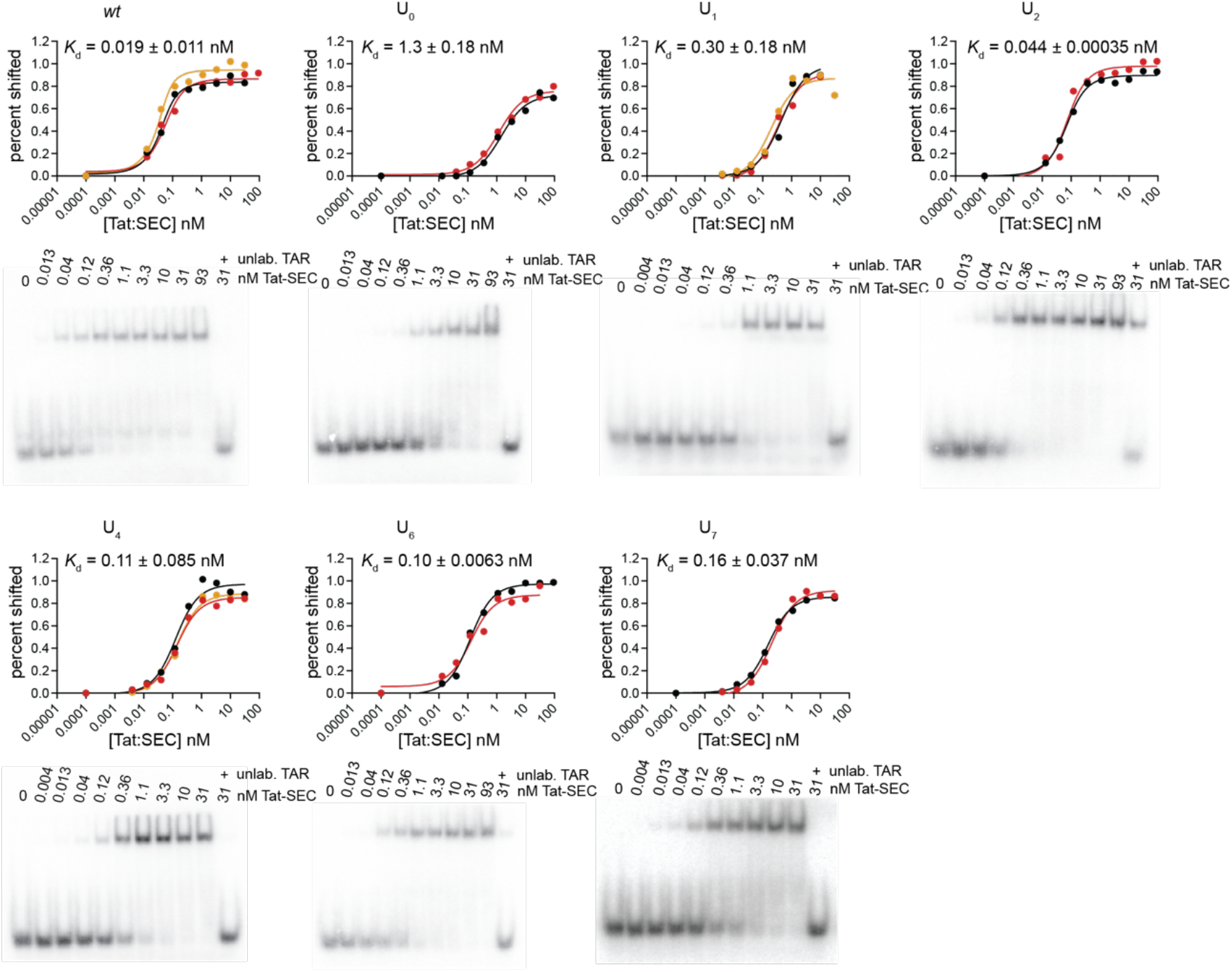
Measurements of TAR-Tat:SEC binding using electrophoretic mobility shift assay (EMSA). Shown are EMSA binding curves for TAR bulge mutants U0,1,2,4,6,7 and UCU along with average apparent *K*_d_ values (see Methods) for each variant, obtained by fitting data to equation 2 using GraphPad Prism (version 9.3.1). Replicate binding curves are overlayed (black: replicate 1, red: replicate 2, orange: replicate 3). Below binding curves for each variant is a representative EMSA gel (replicate 1).

