## Supplementary Materials for "RNA conformational propensities determine cellular activity"

**Supplementary Discussion 1: Mutant behavior that deviates from the model**

The natural variant, $U_2$, deviated from the model, exhibiting binding affinity in the Tat-ARM peptide binding assay (Fig. 2d) that was lower than predicted from its stacking energy (Fig. 2c). The $U_2$ mutant has the shortest bulge length capable of supporting a base-triple[22,25,26], and its topological constraints may result in a smaller proportion of the stacked conformational states having the inter-helical orientation needed to form the base triple[29]. Indeed, analysis of the FARFAR-NMR $U_2$ ensemble[11] reveals that fewer of the stacked conformations are base-triple competent relative to *wt* (Supplementary Table 5). These observations suggest that a more granular description of the stacked-state ensemble may be required to quantify $\Delta\Delta G_{stack}$ for $U_2$ and highlights the power of obtaining new insights from data deviating from predictions.

The $U_7$ variant also displayed a difference in relative free energy change between the Tat-ARM peptide binding assay (Fig. 2d) and the cell-based assay (Fig. 3b). In the peptide assay, $U_7$ binds with higher affinity than $U_6$, breaking the trend of decreasing affinity with bulge length. However, in the cell-based assay and the SEC-binding assay, $U_7$ had a lower level of transactivation relative to $U_6$, maintaining the trend of decreased activity with increasing bulge length. As noted in the main text, there is NMR evidence to suggest that the $U_7$ uridine-rich bulge may be forming an additional helix consisting of U-U mismatches that improves stacking and does not sterically hinder peptide binding. Our observation of several imino resonances in the region expected for U-U mismatches supports this hypothesis (Extended Data Fig. 2). However, in the Tat-SEC complex this stacking benefit may be counteracted by steric interactions, which could account for the decrease in affinity of $U_7$ relative to $U_6$ in the Tat:SEC binding assay and cell-based assay. Indeed, structural modelling revealed steric overlap between the bulge and Cyclin T1 for $U_7$ variant that are not present in models for *wt* or $U_2$ TAR (Supplementary Figure 1).

**Supplementary Discussion 2: Effect of base-triple destabilizing mutations on Tat-ARM peptide binding and cellular transactivation.**

A model that can account for the similar binding affinities measured for Tat-ARM to the base-triple destabilizing TAR mutants is that Tat-ARM binds these TAR mutants in a kinked rather than the stacked conformation (Extended Data Fig. 5a-b). A prior study provided evidence that the Tat-ARM ligand-mimic argininamide can bind TAR in a kinked conformation[7]. If the conformational penalty due to the base-destabilizing mutation ($\Delta G_{triple}$) exceeds the differences between how well Tat-ARM binds the stacked versus kinked conformation ($\Delta\Delta G = \Delta G_{bind,stack} - \Delta G_{bind,kinked}$), then the Tat-ARM peptide would predominantly bind to TAR in the kinked conformation (Extended Data Fig. 5b). Moreover, since the kinked state predominates for all bulge lengths, binding to the kinked conformation would be independent of bulge-length, as observed in Fig. 2d. This model is supported by the lower maximum fluorescence value with saturating Tat-ARM peptide for all the bulge mutants (Extended Data Fig. 5a). Because the fluorescence value is dependent on the orientation of the C and N terminal ends of the peptide relative to one another, this difference suggests that those variants with a lower maximum fluorescence are binding the peptide in a different conformation in which the peptide termini are closer together than when in the base-triple formation.

In contrast, the behavior observed in the cellular context is as predicted from our thermodynamic model which assumes that Tat:SEC binds to the TAR variants predominantly in a stacked base-triple like conformation so that the differences between the binding energetics for the *wt* and disrupted base triple variants is a constant ($c_{\text{triple}}$) across all bulge lengths. In this case, it appears that the conformational penalty due to the base-destabilizing mutation ($\Delta G_{\text{triple}}$) does not exceed the difference between how well Tat:SEC binds the stacked versus kinked conformation ($\Delta\Delta G = \Delta G_{\text{bind,stack}} - \Delta G_{\text{bind,kinked}}$). Tat:SEC could have a stronger preference for binding the stacked versus kinked conformation because a stacked conformation may be required to properly orient the TAR apical loop so that it can form key contacts with the cyclin-T1 component of Tat:SEC[15], which is absent from Tat-ARM (Extended Data Fig. 5b-c). One possibility is that Tat:SEC binds to the base-triple disrupting mutants in a stacked TAR conformation in which U23 forms a U23•U27-A38 base-triple-like conformation with U23 forming a single hydrogen bond with U27 but otherwise retaining the overall *wt* TAR conformation in the quinary complex (Extended Data Fig. 5d).

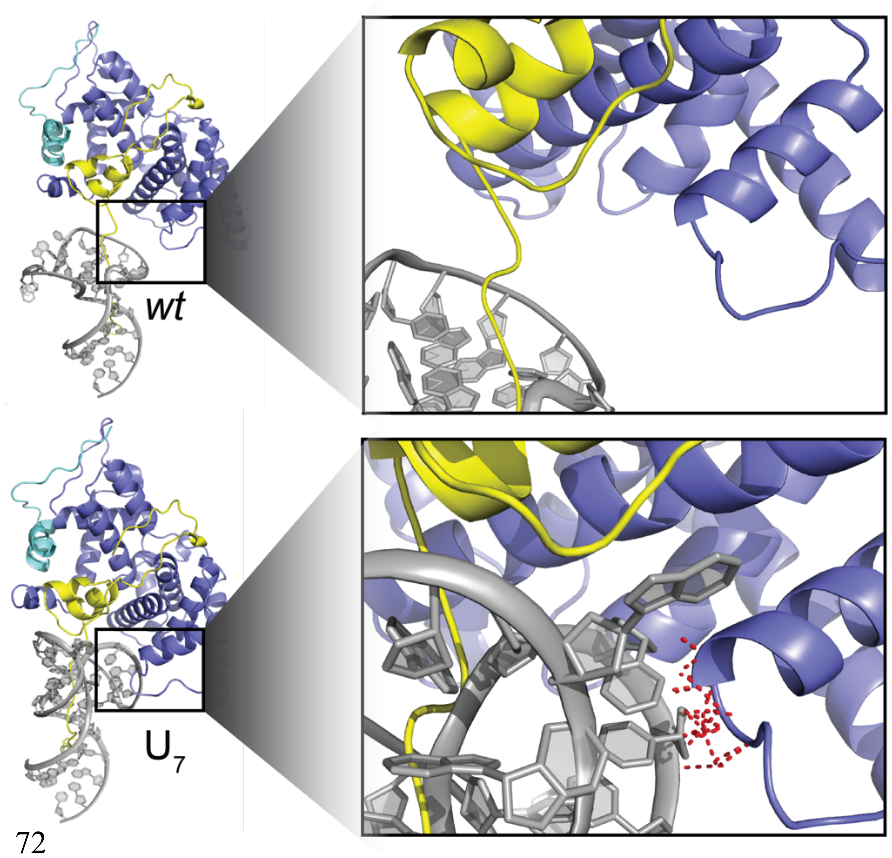


**Supplementary Figure 1. Model of steric interaction between the $U_7$ bulge and P-TEFb.** (Left) FARFAR models of representative base-triple conformations of *wt* and $U_7$ bound to the Tat:SEC complex. (Right) Zoomed in view of the bulge interaction with P-TEFb. In dashed red lines are atom distances between bulge residues and P-TEFb that are within 2.5 Å, representing steric overlap. $U_7$ (bottom) has multiple steric overlaps, whereas *wt* (top) does not.

**Supplementary Table 1. Comparison of the interhelical stacked state population ($p_{stack}$) determined using CSP and FARFAR-NMR.** $p_{stack}$ was determined using these two methods for the *wt* and $U_7$ FARFAR-NMR ensembles[11].

| **RNA** | $p_{stack}$ **CSP** | $p_{stack}$ **FARFAR-NMR** |
|---|---|---|
| *wt* | 0.31 | 0.43 |
| $U_7$ | 0.22 | 0.26 |

**Supplementary Table 2. Measuring stacking energetics using CSPs.** Shown for each TAR variant are the measured chemical shifts (δ) for U23-C6 and A22-C8, the computed population of the stacked state ($p_{stack}$), and the difference in stacking energetics $\Delta\Delta G_{stack}$ referenced to *wt*.

| **RNA** | **$\delta_{U23\text{-}C6}$ (ppm)** | **$\delta_{A22\text{-}C8}$ (ppm)** | **$p_{stack}$ (%)** | **$\Delta\Delta G_{stack}$ (*j-wt*) (kcal/mol)** |
|---|---|---|---|---|
| *wt* | 141.94 | 139.51 | 0.31 | (0) |
| $U_0$ | n/a | 139.07 | n/a | n/a |
| $U_1$ | 143.24 | 139.90 | 0.75 | -1.14 |
| $U_2$ | 143.27 | 139.87 | 0.75 | -1.12 |
| $U_3$ | 142.58 | 139.46 | 0.43 | -0.30 |
| $U_4$ | 141.85 | 139.38 | 0.23 | 0.23 |
| $U_5$ | 141.71 | 139.32 | 0.18 | 0.42 |
| $U_6$ | 141.64 | 139.33 | 0.17 | 0.47 |
| $U_7$ | 141.74 | 139.40 | 0.22 | 0.27 |
| *wt*-A27U/U38A | 141.99 | 139.56 | 0.34 | -0.09 |
| $U_0$-A27U/U38A | n/a | 139.20 | n/a | n/a |
| $U_1$-A27U/U38A | 143.35 | 139.94 | 0.79 | -1.28 |
| $U_2$-A27U/U38A | 143.27 | 140.01 | 0.81 | -1.33 |
| $U_3$-A27U/U38A | 142.50 | 139.43 | 0.40 | -0.23 |
| $U_4$-A27U/U38A | 141.98 | 139.48 | 0.30 | 0.01 |
| $U_5$-A27U/U38A | 141.69 | 139.35 | 0.19 | 0.39 |
| $U_6$-A27U/U38A | 141.68 | 139.33 | 0.18 | 0.43 |
| $U_7$-A27U/U38A | 141.70 | 139.37 | 0.20 | 0.35 |
| *wt*-A27deazaN7 | 141.72 | 139.44 | 0.23 | 0.23 |

| | | | | |
|---|---|---|---|---|
| $U_0$-A27deazaN7 | n/a | 139.04 | n/a | n/a |
| $U_1$-A27deazaN7 | 143.33 | 139.91 | 0.78 | -1.22 |
| $U_2$-A27deazaN7 | 143.15 | 140.01 | 0.78 | -1.23 |
| $U_3$-A27deazaN7 | 142.28 | 139.51 | 0.38 | -0.19 |
| $U_4$-A27deazaN7 | 141.71 | 139.34 | 0.19 | 0.39 |
| $U_5$-A27deazaN7 | 141.54 | 139.26 | 0.12 | 0.72 |
| $U_6$-A27deazaN7 | 141.55 | 139.28 | 0.13 | 0.66 |
| $U_7$-A27deazaN7 | 141.59 | 139.32 | 0.15 | 0.54 |

87

**Supplementary Table 3. Measured $K_d$ and calculated $\Delta G_{pep}$ and $\Delta\Delta G_{pep}(j\text{-}wt)$ values.** Shown for each TAR variant are the measured $K_d$ values and standard deviations for binding to the Tat-ARM peptide, the corresponding $\Delta G_{pep}$ (T= 298.15°K) and standard deviations, and the differences in binding energetics $\Delta\Delta G_{pep}$ referenced to *wt*.

| **RNA** | **Mean $K_d$ (nM)** | **Std. Dev. (nM)** | **Mean $\Delta G_{pep}$ (kcal/mol)** | **Std. Dev. (kcal/mol)** | **Mean $\Delta\Delta G_{mut\text{-}wt}$ (kcal/mol)** | **Std. Dev. (kcal/mol)** |
|---|---|---|---|---|---|---|
| *wt* | 42 | 10 | -9.9 | 0.13 | (0) | (0) |
| $U_0$ | 1100 | 230 | -8.0 | 0.13 | 1.9 | 0.23 |
| $U_1$ | 320 | 50 | -8.7 | 0.10 | 1.7 | 0.17 |
| $U_2$ | 14 | 3 | -10.5 | 0.14 | -0.66 | 0.051 |
| $U_3$ | 33 | 7 | -10.0 | 0.12 | -0.15 | 0.14 |
| $U_4$ | 46 | 2 | -9.8 | 0.026 | 0.056 | 0.14 |
| $U_5$ | 72 | 12 | -9.6 | 0.11 | 0.31 | 0.12 |
| $U_6$ | 100 | 19 | -9.3 | 0.11 | 0.52 | 0.23 |
| $U_7$ | 82 | 15 | -9.5 | 0.11 | 0.39 | 0.15 |
| *wt*-A27U/U38A | 420 | 130 | -8.6 | 0.20 | 1.3 | 0.16 |
| $U_0$-A27U/U38A | 3200 | 1400 | -7.4 | 0.35 | 2.5 | 0.31 |
| $U_1$-A27U/U38A | 610 | 110 | -8.3 | 0.10 | 1.6 | 0.12 |
| $U_2$-A27U/U38A | 300 | 30 | -8.7 | 0.065 | 1.1 | 0.15 |
| $U_3$-A27U/U38A | 380 | 140 | -8.6 | 0.23 | 1.2 | 0.25 |
| $U_4$-A27U/U38A | 300 | 62 | -8.7 | 0.12 | 1.1 | 0.15 |
| $U_5$-A27U/U38A | 340 | 41 | -8.7 | 0.066 | 1.2 | 0.18 |
| $U_6$-A27U/U38A | 220 | 40 | -8.9 | 0.11 | 0.96 | 0.083 |
| $U_7$-A27U/U38A | 180 | 35 | -9.0 | 0.13 | 0.84 | 0.083 |

| | | | | | | |
|---|---|---|---|---|---|---|
| *wt*-A27deazaN7 | 870 | 460 | -8.2 | 0.32 | 1.7 | 0.33 |
| $U_0$-A27deazaN7 | 940 | 330 | -8.1 | 0.18 | 1.8 | 0.22 |
| $U_1$-A27deazaN7 | 630 | 110 | -8.3 | 0.10 | 1.6 | 0.14 |
| $U_2$-A27deazaN7 | 210 | 39 | -8.9 | 0.11 | 0.94 | 0.15 |
| $U_3$-A27deazaN7 | 240 | 160 | -9.0 | 0.50 | 0.86 | 0.43 |
| $U_4$-A27deazaN7 | 340 | 53 | -8.7 | 0.094 | 1.2 | 0.13 |
| $U_5$-A27deazaN7 | 270 | 34 | -8.8 | 0.076 | 1.1 | 0.10 |
| $U_6$-A27deazaN7 | 160 | 36 | -9.1 | 0.14 | 0.77 | 0.24 |
| $U_7$-A27deazaN7 | 110 | 16 | -9.3 | 0.08 | 0.54 | 0.17 |

**Supplementary Table 4. Statistics assessing the quality of the fit for various $\Delta\Delta G$ comparisons.** (Top) Shown are RMSD and $R^2$ values for the fit of each $\Delta\Delta G$ comparison to our thermodynamic model. (Bottom) The RMSD and $R^2$ for the best-fit linear regression, along the parameters of that regression line including slope, y-intercept, and their 95% confidence intervals.

| Model Fit | | | | |
|---|---|---|---|---|
| Parameter | $\Delta\Delta G_{stack}$ *vs.* $\Delta\Delta G_{pep}$ (+$U_2$) | $\Delta\Delta G_{stack}$ *vs.* $\Delta\Delta G_{pep}$ (-$U_2$) | $\Delta\Delta G_{stack}$ *vs.* $\Delta\Delta G_{cell}$ | $\Delta\Delta G_{pep}$ *vs.* $\Delta\Delta G_{cell}$ |
| RMSD (kcal/mol) | 0.24 | 0.18 | 0.56 | 0.41 |
| $R^2$ | 0.62 | 0.58 | -0.54 | 0.71 |

| Best Fit | | | | |
|---|---|---|---|---|
| Parameter | $\Delta\Delta G_{stack}$ *vs.* $\Delta\Delta G_{pep}$ (+$U_2$) | $\Delta\Delta G_{stack}$ *vs.* $\Delta\Delta G_{pep}$ (-$U_2$) | $\Delta\Delta G_{stack}$ *vs.* $\Delta\Delta G_{cell}$ | $\Delta\Delta G_{pep}$ *vs.* $\Delta\Delta G_{cell}$ |
| RMSD (kcal/mol) | 0.17 | 0.17 | 0.37 | 0.40 |
| $R^2$ | 0.83 | 0.61 | 0.27 | 0.74 |
| Slope | 0.68 | 0.81 | 0.43 | 0.94 |
| Slope 95% CI | 0.57 to 0.79 | 0.56 to 1.05 | 0.18 to 0.68 | 0.77 to 1.11 |
| y-intercept (kcal/mol) | 0.07 | 0.04 | 0.24 | 0.16 |
| y-int 95% CI | 0.01 to 0.13 | -0.04 to 0.12 | 0.11 to 0.37 | 0.02 to 0.29 |

**Supplementary Table 5. Analysis of *wt* and $U_2$ FARFAR-NMR ensembles.** The previously determined FARFAR-NMR ensembles[11] were used to compute the population of the stacked ($p_{stack}$) and base-tripled ($p_{triple}$) conformational states for *wt* and $U_2$, along with the corresponding *K*, ΔG, and ΔΔG (referenced to *wt*) values.

| **FARFAR-NMR stacking analysis** | | | | |
|---|---|---|---|---|
| Model | $p_{stack}$ | $K_{stack}$ | $\Delta G_{stack}$ (kcal/mol) | $\Delta\Delta G_{stack}$ ($U_2$-*wt*) (kcal/mol) |
| *wt* | 0.42 | 0.72 | 0.19 | (0) |
| $U_2$ | 0.99 | 99 | -2.7 | -2.9 |
| **FARFAR-NMR base-triple analysis** | | | | |
| Model | $p_{triple}$ | $K_{triple}$ | $\Delta G_{triple}$ (kcal/mol) | $\Delta\Delta G_{triple}$ ($U_2$-*wt*) (kcal/mol) |
| *wt* | 0.03 | 0.031 | 2.1 | (0) |
| $U_2$ | 0.11 | 0.12 | 1.2 | -0.82 |

**Supplementary Table 6. Measured luminescence values and calculated Tat-dependent $\Delta G_{cell}$ and $\Delta\Delta G_{cell}$ (*j-wt*) values:** Shown are the measured FLuc/RLuc luminescence ratios in the presence and absence of the co-transfected Tat plasmid, the calculated Tat-dependent $\Delta G_{cell}$ value (using the incubator T = 37°C / 310.15°K) and their standard deviations, as well as the differences in Tat-dependent transactivation energies $\Delta\Delta G_{cell}$ referenced to *wt* and their standard deviations.

| RNA | Mean FLuc/RLuc + Tat (RLU) | Mean FLuc/RLuc - Tat (RLU) | Mean Tat-dependent $\Delta G_{cell}$ (kcal/mol) | Std. Dev. (kcal/mol) | Mean $\Delta\Delta G_{cell}$ (kcal/mol) | Std. Dev. (kcal/mol) |
|---|---|---|---|---|---|---|
| *wt* | $1.0 \times 10^{-1}$ | $5.9 \times 10^{-4}$ | -2.9 | 0.68 | (0) | (0) |
| $U_0$ | $8.7 \times 10^{-4}$ | $3.4 \times 10^{-4}$ | -0.6 | 0.20 | 2.4 | 0.23 |
| $U_1$ | $5.6 \times 10^{-2}$ | $6.8 \times 10^{-4}$ | -2.4 | 0.87 | 0.69 | 0.36 |
| $U_2$ | $7.5 \times 10^{-2}$ | $5.1 \times 10^{-4}$ | -3.0 | 0.31 | -0.091 | 0.17 |
| $U_3$ | $8.4 \times 10^{-2}$ | $6.0 \times 10^{-4}$ | -3.1 | 0.25 | -0.12 | 0.16 |
| $U_4$ | $1.1 \times 10^{-1}$ | $6.4 \times 10^{-4}$ | -3.0 | 0.31 | 0.067 | 0.15 |
| $U_5$ | $9.1 \times 10^{-2}$ | $5.4 \times 10^{-4}$ | -3.0 | 0.42 | 0.17 | 0.097 |
| $U_6$ | $4.0 \times 10^{-2}$ | $5.1 \times 10^{-4}$ | -2.5 | 0.40 | 0.62 | 0.24 |
| $U_7$ | $2.2 \times 10^{-2}$ | $5.2 \times 10^{-4}$ | -2.1 | 0.45 | 1.0 | 0.23 |
| *wt*-A27U/U38A | $2.4 \times 10^{-2}$ | $6.5 \times 10^{-4}$ | -2.1 | 0.42 | 1.0 | 0.24 |
| $U_0$-A27U/U38A | $3.7 \times 10^{-3}$ | $5.8 \times 10^{-4}$ | -1.1 | 0.22 | 2.0 | 0.28 |
| $U_1$-A27U/U38A | $3.0 \times 10^{-2}$ | $5.7 \times 10^{-4}$ | -2.1 | 0.50 | 1.0 | 0.15 |
| $U_2$-A27U/U38A | $4.0 \times 10^{-2}$ | $7.2 \times 10^{-4}$ | -2.3 | 0.47 | 0.83 | 0.27 |
| $U_3$-A27U/U38A | $1.6 \times 10^{-2}$ | $6.1 \times 10^{-4}$ | -2.0 | 0.24 | 0.94 | 0.21 |
| $U_4$-A27U/U38A | $9.5 \times 10^{-3}$ | $5.1 \times 10^{-4}$ | -1.8 | 0.25 | 1.2 | 0.077 |
| $U_5$-A27U/U38A | $4.1 \times 10^{-3}$ | $4.4 \times 10^{-4}$ | -1.3 | 0.16 | 1.6 | 0.18 |

| | | | | | | |
|---|---|---|---|---|---|---|
| $U_6$-A27U/U38A | 1.8 x $10^{-3}$ | 4.5 x $10^{-4}$ | -0.85 | 0.094 | 2.1 | 0.19 |
| $U_7$-A27U/U38A | 1.4 x $10^{-3}$ | 6.7 x $10^{-4}$ | -0.28 | 0.58 | 2.6 | 0.26 |

111

**Supplementary Table 7. Measured apparent $K_d$ values and corresponding $\Delta G_{prot}$ and $\Delta\Delta G_{prot}$(*j-wt*) values.** Shown for each TAR variant are the apparent measured $K_d$ values and standard deviations for binding to the Tat:SEC protein complex, the corresponding $\Delta G_{prot}$ (T= 298.15°K) and standard deviation, and the differences in binding energetics $\Delta\Delta G_{prot}$ referenced to *wt*.

| RNA | Mean $K_d$ (nM) | Std. Dev. (nM) | Mean $\Delta G_{prot}$ (kcal/mol) | Std. Dev. (kcal/mol) | Mean $\Delta\Delta G_{prot}$ (kcal/mol) | Std. Dev. (kcal/mol) |
|---|---|---|---|---|---|---|
| *wt* | 0.019 | 0.011 | -14.4 | 0.4 | (0) | 0.35 |
| $U_0$ | 1.3 | 0.18 | -11.9 | 0.1 | 2.5 | 0.08 |
| $U_1$ | 0.30 | 0.12 | -12.8 | 0.3 | 1.6 | 0.26 |
| $U_2$ | 0.044 | 0.0004 | -13.8 | 0.005 | 0.57 | 0.005 |
| $U_4$ | 0.11 | 0.0085 | -13.3 | 0.05 | 1.1 | 0.045 |
| $U_6$ | 0.10 | 0.0063 | -13.4 | 0.04 | 1.05 | 0.036 |
| $U_7$ | 0.16 | 0.037 | -13.1 | 0.14 | 1.3 | 0.14 |

**Supplementary Table 8. RNA sequences and oligonucleotides.** This excel spreadsheet contains in tab 1 the sequences of all synthesized RNAs in this study; in tab 2 the sequences of the DNA oligo primers used to construct the TAR-FLuc-140 plasmid; in tab 3 the sequences of the inserted oligos used to create the TAR mutant plasmids used in the cellular transactivation assay.
